# Transgenic mice expressing tunable levels of DUX4 develop characteristic facioscapulohumeral muscular dystrophy-like pathophysiology ranging in severity

**DOI:** 10.1101/471094

**Authors:** Takako I. Jones, Guo-Liang Chew, Pamela Barraza-Flores, Spencer Schreier, Monique Ramirez, Ryan D. Wuebbles, Dean J. Burkin, Robert K. Bradley, Peter L. Jones

## Abstract

**Background:** All types of facioscapulohumeral muscular dystrophy (FSHD) are caused by the aberrant myogenic activation of the somatically silent DUX4 gene, which initiates a cascade of cellular events ultimately leading to FSHD pathophysiology. Therefore, FSHD is a dominant gain-of-function disease that is amenable to modeling by *DUX4* overexpression. However, there is large variability in the patient population. Typically, progressive skeletal muscle weakness becomes noticeable in the second or third decade of life, yet there are many genetically FSHD individuals who develop symptoms much later in life or remain relatively asymptomatic throughout their lives. Conversely, in rare cases, FSHD may present clinically prior to 5-10 yrs of age, ultimately manifesting as a very severe early onset form of the disease. Thus, there is a need to control the timing and severity of pathology in FSHD-like models.

**Methods:** We have recently described a line of conditional *DUX4* transgenic mice, *FLExDUX4*, that develop a myopathy upon induction of human *DUX4-fl* expression in skeletal muscle. Here, we use the *FLExDUX4* mouse crossed with the skeletal muscle-specific and tamoxifen inducible line *ACTA1-MerCreMer* to generate a highly versatile bi-transgenic mouse model with chronic, low-level DUX4-fl expression and mild pathology, that can be induced to develop more severe FSHD-like pathology in a dose-dependent response to tamoxifen. We identified conditions to reproducibly generate models exhibiting mild, moderate, or severe DUX4-dependent pathophysiology, and characterized their progression.

**Results:** We assayed DUX4-fl mRNA and protein levels, fitness, strength, global gene expression, histopathology, and immune response, all of which are consistent with an FSHD-like myopathic phenotype. Importantly, we identified sex-specific and muscle-specific differences that should be considered when using these models for preclinical studies.

**Conclusions:** The *ACTA1-MCM;FLExDUX4* bi-transgenic mouse model expresses a chronic low level of DUX4-fl and has mild pathology and detectable muscle weakness. The onset and progression of moderate to severe pathology can be controlled via tamoxifen injection to provide consistent and readily screenable phenotypes for assessing therapies targeting DUX4-fl mRNA and protein. Thus, these FSHD-like mouse models can be used to study a range of DUX4-fl expression and pathology dependent upon investigator need, through controlled mosaic expression of *DUX4*.

## Introduction

Facioscapulohumeral muscular dystrophy (FSHD) afflicts females and males of all ages and an estimated 1:8 300–15,000 people world-wide [1–4]. All forms of FSHD share a common pathogenic mechanism, increased somatic expression of the DUX4 (Double homeobox 4) retrogene caused by the loss of stable epigenetic repression of the chromosome 4q35.2 D4Z4 macrosatellite array [5–10]. Epigenetic dysregulation of the locus is caused by large deletions of the D4Z4 array on a single 4q allele, reducing it to 1-10 repeat units (RU) (classified as FSHD1) [6, 11, 12], or by mutations in genes encoding repressive epigenetic regulators of the locus (classified as FSHD2) [13, 14]. In addition, all forms of FSHD require a permissive *DUX4* polyadenylation signal (PAS) in *cis* distal to a dysregulated chromosome 4q D4Z4 array [15]. Pathology ultimately results from the aberrant increase in stable somatic expression of the dysregulated *DUX4* mRNA from the distal-most RU [5, 8–10, 15–18].

The *DUX4* gene encodes several alternative mRNA isoforms generated by alternate 5’splice site usage in the first exon [8]; however, only the DUX4-full length (*DUX4-fl*) mRNA is pathogenic when expressed in muscle [8, 15, 19]. *DUX4-fl* encodes a paired homeobox domain transcription factor (DUX4-FL) normally expressed in healthy human testis, pluripotent cells, and cleavage-stage embryos, and the DUX4-mediated transcriptional program is key for zygotic genome activation, all of which supports an important role for DUX4-FL during early embryonic development [5, 8, 20–23]. Adult somatic cells from healthy individuals are typically devoid of detectable *DUX4-fl* expression [8, 15, 16]. However, individuals meeting the genetic criteria for FSHD express stable DUX4-fl mRNA and protein in their skeletal muscles, which aberrantly activates an embryonic gene expression profile [8, 15, 16, 24, 25], ultimately leading to FSHD pathophysiology. Interestingly, low somatic expression of *DUX4-fl* mRNA *per se* is not necessarily pathogenic since expression can be detected in some rare cultures of myogenic cells and muscle biopsies from healthy and asymptomatic FSHD subjects, albeit at levels significantly lower than in equivalent cells and tissues from FSHD-affected subjects [9, 16, 17]. Epigenetic analysis of the pathogenic 4q D4Z4 RU shows that the stability of *DUX4* epigenetic repression correlates with disease presentation among healthy, FSHD1-affected, and FSHD1-asymptomatic subjects [9]. Together, this data supports that the level of somatic *DUX4-fl* expression, which is inducible and affected by the epigenetic stability in the region, is the key determinant of disease onset and severity.

In addition to the complex genetic and epigenetic conditions that are required to develop clinical FSHD, the pathogenic mechanism is also unusual among neuromuscular diseases. While FSHD is an autosomal dominant gain-of-function disease, the pathogenic DUX4 gene is typically expressed in only a small fraction (<1%) of myocytes, ultimately leading to debilitating muscle pathology over time [8, 16, 26]. This may account in part for the more common adult onset of clinical symptoms in FSHD patients [27, 28]. In addition, it appears that FSHD pathology is caused by sporadic bursts of increased DUX4-fl expression in differentiated myocytes, which are epigenetically suppressed in healthy and asymptomatic subjects [9, 19, 29]. Since these cells are syncytial, the detrimental effects of aberrant DUX4-FL expression may be found throughout an FSHD myofiber despite expression initially being restricted to a small percentage of myonuclei at any one time [26]. Regardless, DUX4-fl expression in FSHD myocytes and skeletal muscle, even when bursting, is still extremely rare, highly variable, and difficult to detect [8, 16, 19].

Since DUX4-FL is a transcription factor not typically expressed in healthy muscle, its aberrant expression alters the mRNA profiles of numerous genes in myocytes [24, 30, 31]. Many of these DUX4 target genes, including germline specific genes, cleavage stage genes, and immune system regulators, are not normally expressed in healthy myogenic cells [22, 24, 30, 31]. In addition, DUX4-FL expression ultimately initiates a cascade of numerous potentially detrimental events including the disruption of proteostasis, nonsense-mediated decay, and mRNA splicing mechanisms [19, 32, 33], altered myogenesis [34, 35], and induction of apoptosis [19, 36–38]. These DUX4-mediated changes, either alone or in combination, lead to progressive muscle cell death and ultimately pathology [10, 18]. Thus, the DUX4-fl mRNA and protein are prime targets for therapeutic intervention, and animal models for FSHD should be based on DUX4-fl expression in adult skeletal muscle.

We have previously reported the generation of the *FLExDUX4 (FLExD)* conditional *DUX4-fl* transgenic line of mice, which contains a DUX4 transgene engineered into the *Rosa26* locus using the FLEx directional switch system [39, 40] to bypass the embryonic lethality from leaky embryonic expression of this transgene [41]. Upon Cre-mediated induction, the transgene recombines to express DUX4-fl under control of the *Rosa26* gene promoter (Figure 1A). Importantly, since many elements within *DUX4* are prime targets for sequence-based therapies [42–45], the DUX4 transgene maintains the exon/intron structure of the endogenous human gene, including the 5’ untranslated region (UTR), the endogenous PAS, and the distal auxiliary elements (DAEs) that enhance *DUX4* mRNA cleavage and polyadenylation events [5, 8, 45–47]. Here we show that the *FLExD* mouse model, when crossed with an appropriate inducible Cre mouse, produces a bi-transgenic model with chronic low levels of *DUX4-fl* expression which can be induced at the desired time to develop a more severe pathology, reproducibly recapitulating many aspects of FSHD pathophysiology. Importantly, the severity of pathology can be controlled by the investigator, thus providing several suitable models for therapeutic interventions targeting DUX4-fl mRNA, protein, and certain downstream pathways.

**Figure 1:**
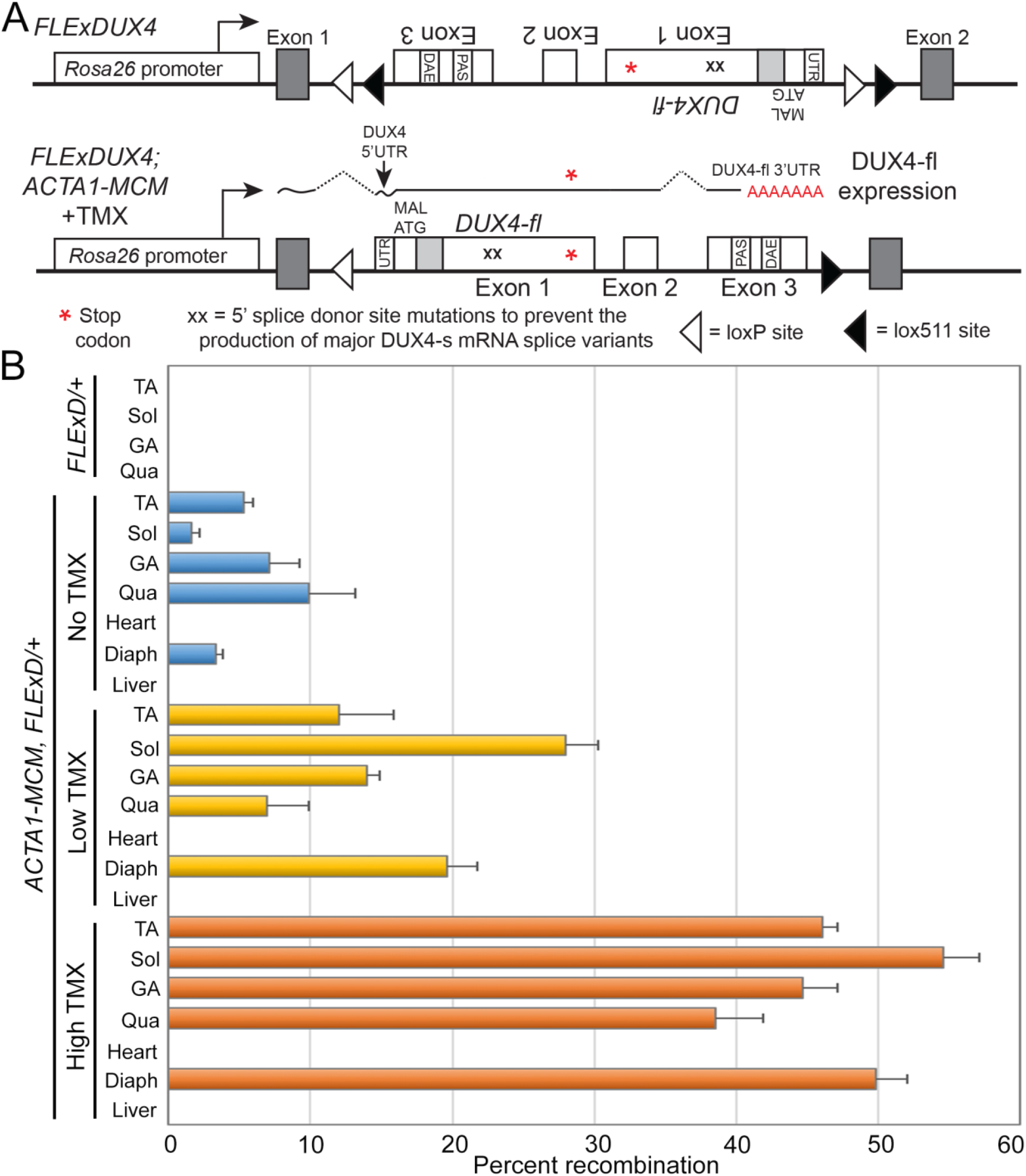
Transgene map and FSHD-like model generation. A) The synthesized FLExDUX4 transgene, flanked by heterologous lox sites (loxP and lox511), was inserted in the antisense orientation to the *Rosa26* promoter and maintains the intron/exon structure and *cis* mRNA regulatory including features, including the PAS and DAE, of the human chromosome 4q35 *DUX4* gene. When crossed with *ACTA1-MCM* mice, the bi-transgenic offspring have the capacity for dosage dependent, TMX-inducible unidirectional recombination of the transgene, resulting in DUX4 expression exclusively in skeletal muscle and transcribed from the *Rosa26* promoter, processed, and terminated in exon 3 using the *DUX4* PAS. B) Genomic PCR indicating percent transgene recombination in different muscles from *FLExD/+* mice and bi-transgenic *ACTA1-MCM;FLExD* mice with no TMX or 3 days after a single IP injection of 5mg/kg TMX (low), or two IP injections of 10mg/kg TMX (high). TA: tibalis anterior, Sol: soleus, GA: gastrocnemius, Qua: quadriceps, Diaph: diaphragm.

## Methods

### Transgenic mice

The *ACTA1-MCM* Cre driver line refers to B6.Cg-Tg(ACTA1-cre/Esr1*)2Kesr/J (JAX 025750) [48] and Rosa26^NZG^ refers to FVB.Cg-Gt(ROSA)26Sor<tm1(CAG-lacZ,-EGFP)Glh>/J (JAX 012429) [49], both purchased from Jackson Labs (Bangor, Maine). *FLExDUX4* (*FLExD*) mice were generated and characterized by the PL Jones lab [41] and are available from Jackson Labs (JAX 028710). Mice were genotyped as described [41].

### Tamoxifen (TMX) injections

Tamoxifen free base (Sigma T5648) was dissolved in 100% ethanol (200mg/ml) at 55°C and added to pre-warmed sterile corn oil (ThermoFisher S25271) to make 20mg/ml frozen stocks. Stock TMX aliquots were warmed to 37°C, diluted with pre-warmed sterile corn oil either 10- or 20-fold just prior to use, and administered to mice by intraperitoneal (IP) injection at a final concentration of 5 or 10mg/kg. For the moderate model, *ACTA1-MCM;FLExD* bi-transgenic animals were injected once with 5mg/kg TMX. For the severe model, *ACTA1-MCM;FLExD* bi-transgenic mice were injected on two consecutive days with 10mg/kg TMX.

### X-Gal staining for evaluating TMX dosing

Male *ACTA1-MCM* mice were crossed with female *Rosa26^NZG^* mice to assess induction of Cre-mediated recombination using our TMX dosage regimens. Control or bi-transgenic mice were treated with TMX, as above. Whole gastrocnemius muscles were fixed in LacZ fixative (0.2% glutaraldehyde, 5mM EGTA, 100mM MgCl_2_ in PBS pH7.4) at 4°C for 4 hours, cryopreserved in cold 30% sucrose/PBS at 4°C overnight, then frozen in O.C.T. for sectioning. Cryosections (8μm) were fixed with 4% paraformaldehyde (PFA)/PBS, then stained with X-Gal staining solution (1X PBS with 2mM MgCl_2_, 5mM potassium ferricyanide, 5mM potassium ferrocyanide, 0.01% Nonidet P-40, 0.1% sodium deoxycholate, 1mg/mL X-Gal) for 50 min. Stained sections were post-fixed in buffered formalin at 4°C for 5 min, rinsed three times with PBS and then deionized water before mounting. Images were captured using an Olympus CX41 microscope with PixeLINK camera under bright field.

### Gene expression analysis by RT-PCR

Total RNA was extracted from dissected mouse muscles homogenized in 10 volumes of TRIzol (ThermoFisher) using the TissueLyser LT (Qiagen), as per manufacturer’s instructions followed by on-column DNaseI treatment and clean-up using the RNeasy mini kit (Qiagen). Quantitative *DUX4-fl* mRNA expression was analyzed using nested qRT-PCR, as described [41, 50]. Expression of DUX4 target genes was analyzed by qPCR using 5 to 10ng cDNA, as described [30, 41]. All oligonucleotide primer sequences for *DUX4-fl*, downstream targets, and *18S* rRNA are previously reported [29]. Expression of Myostatin mRNA was analyzed by qRT-PCR using primers 5’-AGTGGATCTAAATGAGGGCAGT-3’ (forward) and 5’-GTTTCCAGGCGCAGCTTAC-3’ (reverse). Significance was calculated using the two-tailed t test.

### Histology

Freshly dissected muscles were kept moist, coated with O.C.T. compound, snap-frozen in liquid nitrogen-cooled isopentane, and stored at −80°C until sectioning. 12μm cryosections were mounted on slides and air-dried before staining or storage. Sections were used for H&E staining [41] or picrosirius red staining.

### Picrosirius red (SR) staining

SR staining was performed as described [51]. Cryosections (12μm) cut mid belly of the tibialis anterior muscle were fixed with 4% PFA/PBS, pH 7.4 for 10 min, rinsed with dH2O, and dehydrated with a series of 1 min ethanol washes (70%, 95%, 100%) and air dried. Sections were then stained for 1 hr in SR solution (0.1% direct red 80, 1.3% saturated picric acid), and washed three times with dH2O. Stained sections were dehydrated with a series of 1 min ethanol washes (70%, 95%, and 100%), cleared with xylene for 5 min, and mounted. A series of micrographs from each muscle section were captured using a 10X objective on a Leica DM2000 and reconstituted to form a whole muscle cross-section using LAS 4.12 software (Leica Microsystems, Inc.). Whole cross-section images were divided into 3-5 sections and processed with MATLAB (Mathworks) to determine the number of pixels stained red and the total number of pixels stained. Muscles from 3-6 mice for each treatment were analyzed. Significance was calculated by one-way ANOVA using Prism 7 (Graphpad).

### Immunofluorescence (IF)

The 10 μm cryosections of muscle tissue were fixed with 4% PFA/PBS on ice for 10 min, permeabilized with 0.25% TritonX-100/PBS for 10 min, then incubated with blocking solution (5% normal goat serum, 2% BSA, 0.01% TritonX-100/PBS) for 30 min, and then incubated with primary antibody at 4°C overnight. The following day, sections were incubated with secondary antibody at room temperature for 40 min, then mounted in ProLong Gold with DAPI for nuclear staining. DUX4 was detected using E5-5 antibody (Abcam ab124699) diluted 1:200. Embryonic fast myosin heavy chain was detected using the F1.652 monoclonal antibody, developed by the Baxter Lab for Stem Cell Biology, Stanford University, and obtained from the Developmental Studies Hybridoma Bank, created by the NICHD of the NIH and maintained at The University of Iowa, Department of Biology, Iowa City, IA 52242. Dystrophin was detected using an anti-dystrophin rabbit polyclonal antibody (Abcam, ab15277). Immunofluorescent images were captured using the Leica DMi8, DFC365 FX camera and LAS X Expert software (Leica Microsystems Inc.).

### Apoptosis Assay

Apoptotic events were analyzed by TUNEL (terminal deoxynucleotide transferase dUTP nick end labeling) staining using the *In Situ* Cell Death Fluorescein Kit (Roche/SIGMA 11684795910). The 10-μm cryosections of TA muscles were mounted on slides and fixed with 4% paraformaldehyde for 20 min. Staining was performed as per manufacturer’s instructions.

### Treadmill exhaustion test

All treadmill tests were performed with an Exer3/6 treadmill and shock detection system (Columbus Instruments) in the mouse mode with electric shocking grid. All mice were acclimated to the apparatus before running by placing them on an unmoving treadmill for 5 min, then at a speed of 5 m/min for 5 min at 0° incline. Mice were rested for 2 days before the first test. The exhaustion test was established for these FSHD-like model mice after several modifications of the Treat-NMD protocol DMD_M.2.1.003. The test was performed using a 7° incline and an initial speed of 5 m/min with speed increasing by 0.5 m every minute. Mice were run until they were unable to maintain a speed to remain off the shock grid for more than 5 seconds (time of fatigue) or a maximum of 20 min (approximate maximum speed is 15 m/min and maximum distance is 200 m). This testing was performed at least three times per mouse with at least two days of rest in between tests. Significance was calculated by two-way ANOVA using Prism 7 (Graphpad).

### *Ex vivo* muscle contractile properties

EDL muscles were excised from deeply anesthetized mice with 3% isoflurane, hung from a computer-controlled servomotor (300B, Aurora Scientific, Inc), and mounted in a heated (30°C) oxygenated tissue bath containing a physiologic salt solution as described [52]. Experiments were performed and data were analyzed using DMA software (Aurora Scientific, Inc) and Prism 7 (Graphpad), as previously described (Treat-NMD SOP, DMD_M.1.2.002 and [53]). Significance for twitch and tetanus was calculated by one-way ANOVA, and for the force frequency analysis it was calculated by two-way ANOVA using Prism 7 (GraphPad).

### RNA-seq

RNA-seq was performed by Genewiz LLC (South Plainfield, NJ). Total RNA (5μg) was isolated from gastrocnemious muscles of 3 mice per group (*ACTA1-MCM, FLExD, ACTA1-MCM;FLExD* moderate TMX, and *ACTA1-MCM;FLExD* severe TMX), as described above. mRNAs were purified using poly(A) selection and then fragmented. First strand cDNA synthesis used random priming followed by second strand synthesis. The resulting double strand cDNA was end repaired, phosphorylated, and A-tailed. Adapters were ligated and PCR amplification was performed. The library was sequenced using the Illumina HiSeq2500platform in a 1×50 base pair single-read configuration in Rapid Run mode, with a total of at least 120 million reads per lane. Sequence reads were trimmed to remove adapter sequences and poor quality nucleotides (error rate <0.05) at the ends. Sequence reads shorter than 50 nucleotides were discarded.

### Differential RNA-seq expression and gene ontology (GO) analysis

#### Data sources

RNA-seq reads for C2C12 expressing DUX4-fl or control were downloaded from the Gene Expression Omnibus (accession numbers GSE87282) [23].

#### Genome annotation, read mapping, and gene expression estimation

Human (hg19) and mouse (mm10) genome annotations were created by merging the UCSC knownGene [54], Ensembl 71 [55], and MISO v2.0 [56] annotations. Sequence reads were mapped to these annotations as previously described [57]. In brief, RSEM v1.2.4 [58] was modified to call Bowtie [59] with the option ‘-v 2’ and then used to map all reads to the merged genome annotation. Remaining unaligned reads were then mapped to the genome and a database of potential splice junctions with TopHat v2.0.8b [60]. All gene expression estimates were normalized using the trimmed mean of M values (TMM) method [61].

Differentially expressed genes (Table S1 and GEO Accession # pending) in DUX4-induced versus control experiments were defined at a threshold of 2-fold change and Bayes factor ≥10 (computed with Wagenmakers’s framework [62]), as observed in all possible pair-wise comparisons of experimental replicates. GO terms that were enriched amongst genes that exhibited increased or decreased expression in DUX4-induced versus control samples were identified with the GOseq method [63], with a false-discovery rate threshold of 0.01. GO superterms (Table S4) are all terms that are descendent of GO terms containing the following key terms, respectively:

<u>Cell cycle</u>: cell cycle, mitosis, mitotic, chromosome segregation, cell division, nuclear division, proliferation, chromosome condensation, kinetochore, spindle, cyclin
<u>Apoptosis</u>: apoptosis, cell death, apoptotic, cell killing
<u>Muscle</u>: muscle, sarco, myofibril
<u>Immune</u>: immune, chemokine, interferon, wound, virus, cytokine, cytokine-, leukocyte, interleukin, Toll-like, toll-like, bacteri, inflamma, defense, immunological, immunoglobulin receptor, MHC, immunoglobulin, viral

Differentially-expressed genes from the comparison of 9 control and 9 FSHD1 muscle biopsies [24] were defined as genes that were consistently differentially expressed at a threshold of Bayes factor ≥10 (computed with Wagenmakers’s framework [62]) in more than 50 out of 81 possible pair-wise comparisons between control and FSHD1 muscles. The intersection of these genes with orthologous differentially-expressed mouse genes from DUX4-induced vs control experiments are summarized in Table S3.

#### Data analysis and visualization

All data analysis was performed in the R programming environment and relied on Bioconductor [64], dplyr [65] and ggplot2 [66]. Venn diagrams were plotted using the venneuler package.

### Alternative splicing analysis

The MISO computational method [56] was used to characterize skipped exon (SE) and retained intron (RI) alternative splicing events for the 12 RNA-seq samples (Tables 3 and S6). MISO (version 0.5.3) was executed with mapping results produced with TopHat, using the following configuration: --read-len:51, min_event_reads = 20, burn_in = 500, lag = 10, num_iters = 5000, and num_chains = 6.

**Table 1:** FSHD-like mouse severity models

| Genotype | TMX | Transgene recombination | DUX4-fl expression | Phenotype |
| --- | --- | --- | --- | --- |
| *ACTA1-MCM/+* | 5mg/kg x 1 IP | NA | No | Control Moderate |
| *ACTA1-MCM/+* | 10mg/kg x 2 IP | NA | No | Control Severe |
| *ACTA1-MCM;FLExD/+* | None | 2-10% | Yes | Mild |
| *ACTA1-MCM;FLExD/+* | 5mg/kg x 1 IP | 7-28% | Yes | Moderate |
| *ACTA1-MCM;FLExD/+* | 10mg/kg x 2 IP | 38-55% | Yes | Severe |

**Table 2:** GO superterms^1^ enriched in FSHD-like models.

| **Comparison** | **Status** | **GO Superterm** | **% of Genes Associated<sup>2</sup>** |
| --- | --- | --- | --- |
| iDUX4 Dox vs vehicle | up | Apoptosis | 16.42 |
| iDUX4 Dox vs vehicle | up | Cell cycle | 12.55 |
| iDUX4 Dox vs vehicle | up | Immune | 0.53 |
| iDUX4 Dox vs vehicle | down | Cell cycle | 39.25 |
| iDUX4 Dox vs vehicle | down | Immune | 7.07 |
| iDUX4 Dox vs vehicle | down | Muscle | 4.99 |
| Moderate model vs MCM | up | Immune | 30.06 |
| Moderate model vs MCM | up | Cell cycle | 24.91 |
| Moderate model vs MCM | up | Apoptosis | 10.29 |
| Moderate model vs MCM | down | Muscle | 8.23 |
| Severe model vs MCM | up | Immune | 31.05 |
| Severe model vs MCM | up | Cell cycle | 26.43 |
| Severe model vs MCM | up | Apoptosis | 16.95 |
| Severe model vs MCM | up | Muscle | 0.70 |
| Severe model vs MCM | down | Muscle | 14.84 |

1 Superterms listed in Table S4
2 Genes enriched for each comparison are listed in Table S5

**Table 3:**
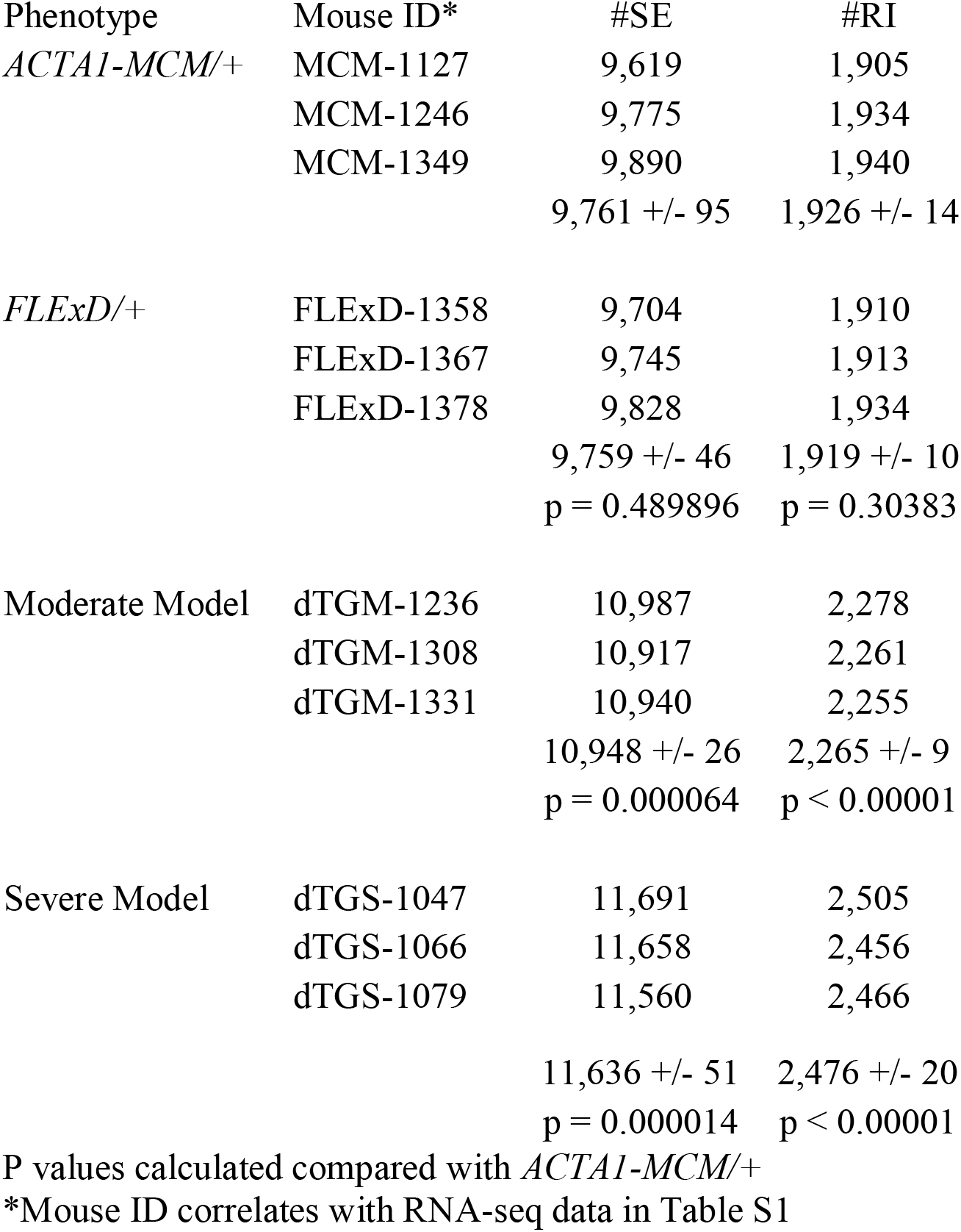
Alternative splicing analysis

## Results

### Generation of mice with mosaic DUX4-fl expression in skeletal muscles

FSHD is caused by mosaic expression of *DUX4-fl* mRNA and its encoded protein from the normally silent *DUX4* gene in a small fraction of differentiated adult skeletal myocytes [8, 19]. Previously we showed that mosaic expression of *DUX4-fl* in skeletal muscle of *FLExDUX4* transgenic mice (or *FLExD)* can produce a very severe myopathy with FSHD-like pathology [41]. However, preclinical testing for different candidate FSHD therapeutics targeting *DUX4-fl* mRNA and protein expression in these mice will likely require different criteria, such as degree and progression of pathophysiology, dependent upon the approach. To address this issue, we generated and characterized a highly reproducible series of phenotypic FSHD-like transgenic mouse models varying in severity and pathogenic progression based on differing levels of mosaic expression of the pathogenic *DUX4-fl* mRNA isoform of human *DUX4* in adult murine skeletal muscle. We identified the tamoxifen (TMX) inducible and skeletal muscle-specific Cre expressing transgenic mice, *ACTA1-MerCreMer* (or *ACTA1-MCM)* [48] as a strong candidate for the generation of the desired phenotypes. To test if these could be used to generate mosaic expression in skeletal muscles and to optimize TMX dosing, the *ACTA1-MCM* mice were crossed with *R26^NZG^* Cre reporter mice [49] that produce readily detectable nuclear ß-galactosidase (*nLacZ*) expression in all cells where Cre is functional in the nucleus (Figure S1). Two dosing regimens of TMX were tested in the *ACTA1-MCM;R26^NZG^* bi-transgenic offspring, a single low dose (5mg/kg) intraperitoneal (IP) injection and two IP injections on consecutive days of a higher dose (10mg/kg). Expression of the *nLacZ* reporter gene was visualized by X-Gal staining [67] in the gastrocnemius muscles isolated two weeks after TMX injection. The results indicated that both dosing regimens produced mosaic patterns of Cre-mediated recombination in skeletal muscle, with the high TMX dose producing ~1.5X more X-Gal stained nuclei than the low dose (Figures S1 and S2). Surprisingly, in the absence of TMX the *ACTA1-MCM;R26^NZG^* mice also showed transgene recombination, although only in a very small fraction of skeletal muscle nuclei (Figure S1C). Interestingly, this low-level recombination and nLacZ expression was not uniform across all nuclei; rather, it also showed a mosaic pattern of recombination (~one-third the number of recombined nuclei induced by the low TMX dose, Figure S2C compared with S2D). Since there was no indication of recombination in the *R26^NZG^* single transgenic animals treated with TMX (Figure S1B), we interpret this result as indicating a low level of MerCreMer protein leaking into the nucleus in the absence of TMX, either at a level that only occasionally leads to recombination or the protein is only sporadically leaky into a few nuclei. Regardless, this suggested that bi-transgenic animals generated with the *ACTA1-MCM* line may be useful for generating very low mosaic expression of a transgene in the absence of TMX induction. Overall, we concluded that the *ACTA1-MCM* line of mice was suitable for generating a range of mosaic transgene expression models.

Generating FSHD-like model mice with differing levels of pathophysiology required adjusting the level of mosaicism with respect to DUX4-fl expression. Therefore, *ACTA1-MCM* mice were crossed with *FLExD* mice [41] (Figure 1A), and 13-14 week old *ACTA1-MCM;FLExD* bi-transgenic mice were treated with the above TMX dosing regimens to induce differing levels of mosaic transgene recombination. Genomic DNA was isolated 3 days post-injection (DPI) of TMX and assayed for transgene recombination by genomic PCR (Figure 1B). The *FLExD* hemizygous mice showed no transgene recombination in the absence of Cre, and the bi-transgenic animals showed variable low levels (2-10%) of transgene recombination in skeletal muscles, but not in heart or liver, in the absence of TMX, due to the above-mentioned sporadic leaky nuclear Cre activity in *ACTA1-MCM* mice. When injected with TMX to induce Cre nuclear activity and stimulate transgene recombination in skeletal muscles, the bi-transgenic mice showed increased recombination (8-30%) in response to the low TMX dose and an even higher level of recombination (38-55%) in response to the high TMX dose. Surprisingly, in all three bi-transgenic models (no, low, and high TMX) there were muscle-specific differences in the transgene recombination rate; however, these differences were consistent between the TMX treated lines (Figure 1B). For example, quadriceps muscles showed the lowest recombination in both TMX-induced models, followed by the tibialis anterior (TA) and gastrocnemius muscles with intermediate levels, while the diaphragm and soleus showed the highest recombination levels. As expected when using the *ACTA1-MCM* driver line, the heart and liver showed no transgene recombination in any of the bi-transgenic animals. Since this assay measures transgene recombination of a single copy transgene per nucleus, the results showing less than 100% recombination represent mosaic recombination which should translate into the desired mosaic DUX4-fl transgene expression. We conclude that we have identified three conditions that reproducibly produce differing levels of mosaic transgene recombination, which we will refer to as mild, moderate, and severe (Table 1), based on the subsequent characterizations described below. Importantly for future studies, we show that different skeletal muscles show different levels of recombination in response to these TMX treatments.

To assess if the variable rates of transgene recombination in each model translated similarly to variable levels of protein expression at the single nucleus level, muscle tissues were analyzed by immunofluorescence (IF) using an anti-DUX4-FL antibody. Mosaic patterns of nuclear DUX4-FL expression were readily detected in cross sections of TA muscle from all three models (Figure 2). The bi-transgenic *ACTA1-MCM;FLExD* mice showed few DUX4-FL positive nuclei in the absence of TMX (Figure 2M and N), consistent with a very low level of transgene recombination (Figure 1B). However, since DUX4-FL is cytotoxic and it is not known how long DUX4-FL expressing cells may remain in the muscles, an IF time course study was performed for the TMX-injected mice. The low-dose mice, which appeared to be moderately affected phenotypically over time and will be referred to as the moderate FSHD-like model (Table 1, discussed in detail below), were assayed over 28 DPI. DUX4-FL expression appeared by 3 DPI (moderate day 3, or MD3), peaking at MD14, and then was greatly reduced by MD28, likely due to the death of DUX4-positive cells (Figure 2A-J). Mice injected with the high-dose TMX regimen were so severely affected by 9 DPI that they had to be sacrificed and could not be assessed further. These mice, which will be referred to as the severe FSHD-like model, similarly showed DUX4-FL expression by three days after the first injection (severe day 3, or SD3) and peak DUX4-FL expression at SD9. Overall, bi-transgenic mice without TMX (referred to as the mild FSHD-like model) exhibited very low numbers of DUX4-FL-positive myonuclei, with the moderate model exhibiting increased numbers, and the severe model the highest numbers. DUX4-FL protein was not detectable in heart or skeletal muscles from the non-recombined *FLExD* mice or *ACTA1-MCM* controls. Thus, each model consistently displays mosaic nuclear DUX4-FL protein expression patterns and provides an indication of the relative abundance of DUX4-FL-expressing nuclei in bi-transgenic animals and in response to two different TMX treatments.

**Figure 2:**
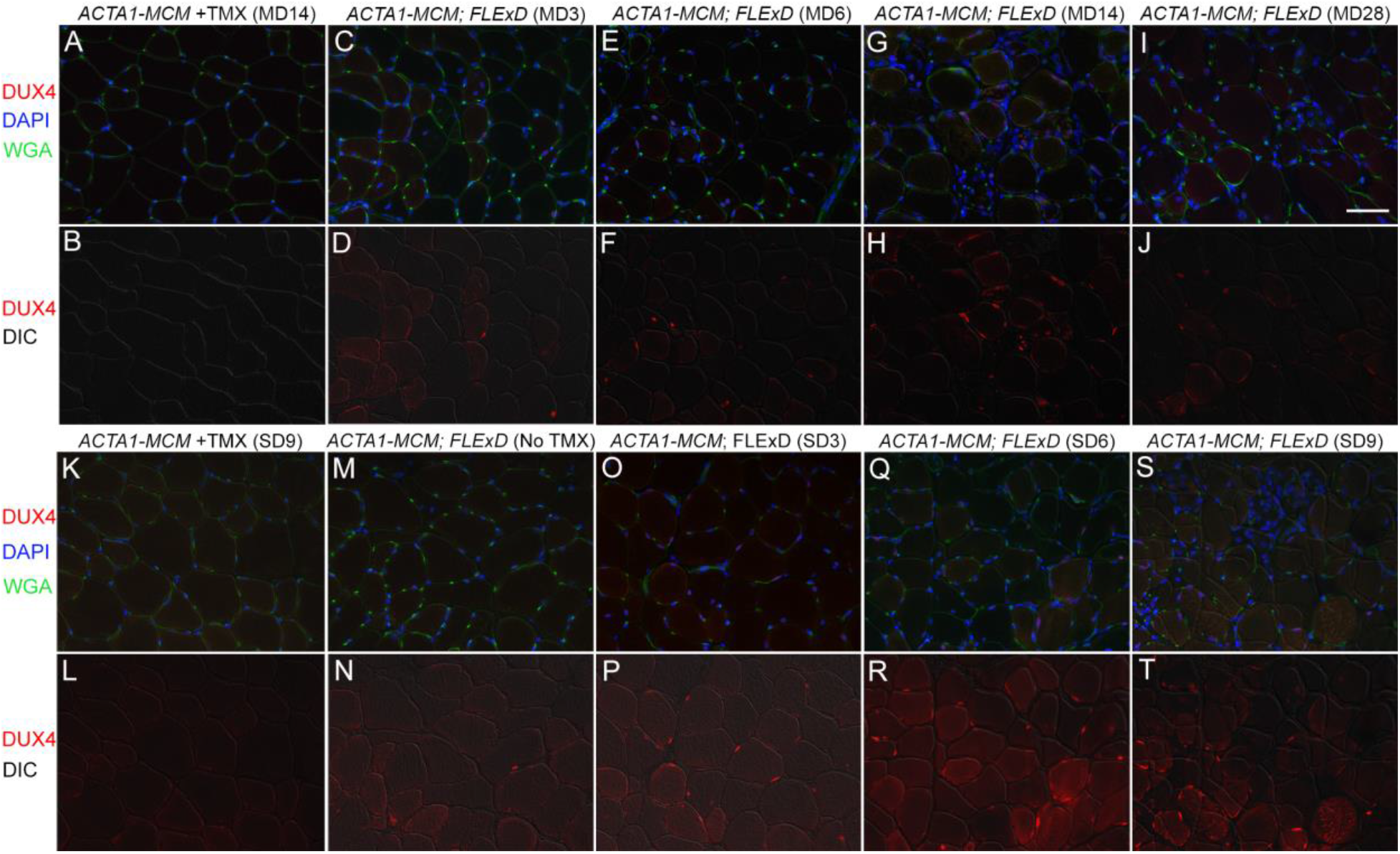
All bi-transgenic *ACTA1-MCM;FLExD* mouse models show mosaic patterns of nuclear DUX4-FL expression. Tibialis anterior muscles were analyzed by IF for DUX4-FL protein expression over 28 days (moderate model, panels C-J) or 9 days (severe model, panels O-T). Muscles from the mild model, bi-transgenic *ACTA1-MCM;FLExD* without TMX (Panels M and N), showed low steady levels of DUX4-FL immunostaining. Muscles from *ACTA1-MCM* mice treated with TMX at MD14 (panels A and B) or SD9 (K and L) showed no DUX4-FL signal and served as negative controls. Red = DUX4 IF, Blue = DAPI staining, Green = WGA staining. Scale bar = 50μm

In order to quantitate the changes in gene expression for each severity model, qRT-PCR was used to measure overall *DUX4-fl* mRNA levels (Figure 3A). However, we have previously shown that this assay is a poor measure of *DUX4-fl* transgene expression using *FLExDUX4* mouse models [41], and *DUX4-fl* mRNA is even difficult to detect in muscle biopsies from FSHD affected subjects [16]. Since DUX4-FL functions as a transcriptional activator in both human and mouse cells [30, 68], expression of DUX4-FL direct target genes has proven to be a more accurate indicator of DUX4-FL expression levels in both species [24, 31, 41]. Therefore, in addition to *DUX4-fl* mRNA, the mRNA levels of two mouse homologs of DUX4-FL direct target genes, *Wfdc3* and *Trim36* [41, 68], were also assayed (Figure 3B and C). Detectable *DUX4-fl* mRNA levels were extremely low in gastrocnemius muscles from all models (Figure 3A), consistent with previous studies [41]. Interestingly, there were no significant changes detected in *DUX4-fl* mRNA levels between the mild, moderate, and severe models 9 days after TMX treatments, a timepoint with prominent differences in DUX4-FL protein expression (Figure 2). In contrast, both DUX4-FL target genes assayed showed significant induction in all bi-transgenic animals compared with the *FLExD/+* mice, indicating the presence of DUX4-FL protein. *Wdfc3* and *Trim36* mRNA levels are each significantly increased in muscles from the moderate and severe models compared with the mild model, and *Trim36* mRNA levels are significantly increased in the severe model compared to the moderate model (Figure 3B and C). We conclude that the bi-transgenic mice, which show increased DUX4-FL expression correlating with the degree of TMX treatment, resemble FSHD patient muscle biopsies, which show increased expression of known DUX4-FL target genes compared with control biopsies [24].

**Figure 3:**
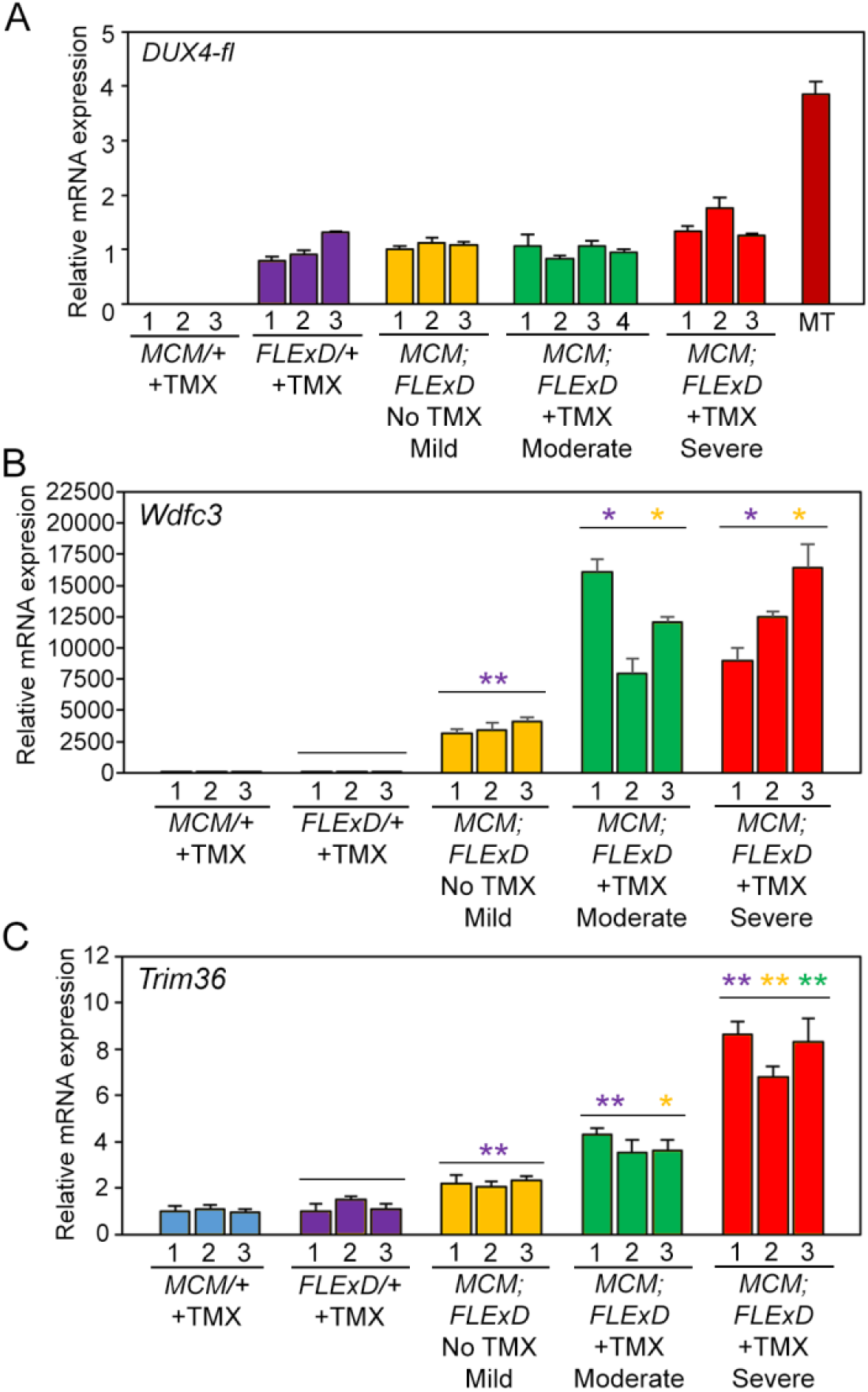
DUX4-FL target genes, but not DUX4-fl mRNA, show TMX dose-dependent increased expression. Gastrocnemius muscles from 13-14 weeks old female mice isolated nine days after TMX administration were assayed for gene expression levels by qRT-PCR. A) Levels of DUX4-fl mRNA are not significantly different between the *FLExDUX4/+* mice and the three bi-transgenic models. However, mRNA expression of B) *Wdfc3* and C) *Trim36* are increased significantly in all bi-transgenic models compared with *FLExDUX4*. Both *Wdfc3* and *Trim36* are significantly increased by moderate and severe TMX induction compared with no TMX. *Trim36* mRNA levels are significantly increased in the severe compared to moderate models. All expression is normalized to Rpl37 expression. MT = equivalent level of cDNA from 17Abic FSHD1 myotubes [16]. Significance calculated with Welch’s t test * p<0.05, ** p<0.01

### Characterization of muscle function for three levels of phenotypic severity

To determine if increases in DUX4-FL protein and target gene expression levels in skeletal muscles correlate with altered muscle function and strength, we performed treadmill exhaustion assays and *ex vivo* muscle physiology studies. Adult *ACTA1-MCM* control and bi-transgenic mice showed size differences between males, with bi-transgenic males being significantly smaller than controls (~26g and ~23g, respectively; Figure S3B and D), while females for both genotypes showed no significant size differences prior to TMX injection (~22g and ~21g, respectively; Figure S3A and C). Therefore, male and female mice were analyzed separately to assess potential sex differences in presentation of the phenotypes. The treadmill analysis protocol consisted of running the mice on a slightly inclined treadmill, slowly increasing the speed until the mice could not run any longer, and measuring the time to fatigue, with a maximum assay time of 20 min. We found these particular conditions (see Methods) provided highly consistent results and a readily assayable window for these three mouse models. Both male and female *ACTA1-MCM* control mice, injected with the appropriate TMX dose for the group being assayed, showed steady levels of near maximum treadmill running fitness (~20 min) over the course of the assays (Figure 4, blue lines). Similarly, treadmill fitness for male and female bi-transgenic mice was not significantly different from the *ACTA1-MCM* controls (Figure 4, green lines). However, male and female mice injected with TMX to produce either a moderate level of pathology (1X 5mg/kg TMX) or a severe level of pathology (2X 10mg/kg TMX) were significantly less fit compared with controls (Figure 4, red lines). Interestingly, both the moderate and severe models showed sex-specific treadmill fitness profiles in which the female mice (Figure 4A, C) were more severely affected than the male mice (Figure 4B, D). Moderate female mice showed significant declines in fitness 10 days after TMX induction (MD10), dropping from near maximum running times to less than 3 minutes. This decline remained at MD16 before beginning to recover to ~4 min running at MD23 and ~10 min by MD29. In comparison, all male moderate model mice showed a significant decline starting at MD6 (running ~15 min) which progressed more slowly than in females, running for ~10 min at MD10 and ~5 min at MD14 and MD17 before recovering to ~15 min by MD28. Overall, although all moderately affected mice showed significant declines in treadmill fitness over the course of four weeks, female mice were more severely affected and recovered more slowly than male mice. For the severe model mice, female mice declined more rapidly than males, although both showed a significant decline in running time before fatigue, decreasing by over 10 min compared with controls by SD5, a point at which the moderate model mice were still unaffected. Female mice could not even begin the assay at SD7, while their male counterparts were slightly less fatigued at SD5 and were able to run a few seconds at SD7. These male mice reached the point of being unable to safely start the assay at SD8. All of these assay points indicated a much more severe phenotype than at any point during the assessment of the moderate model, and these severely affected mice had to be humanely sacrificed by SD10, with general movement in the cage greatly impaired and no signs of recovery. These sex-specific differences make it vitally important to analyze and compare mice of the same sex when performing fitness assays using both of these FSHD-like models.

**Figure 4:**
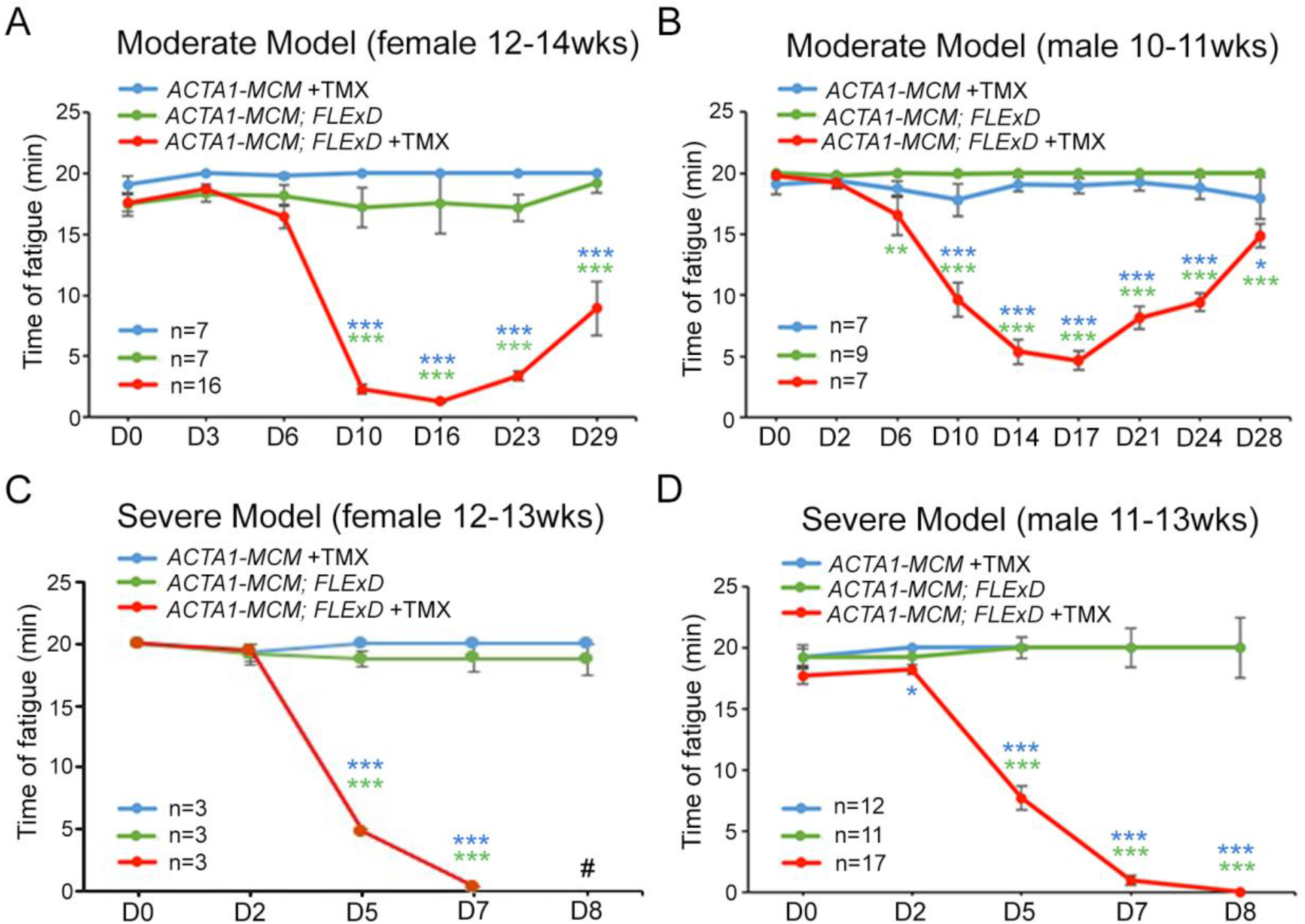
The moderate and severe FSHD-like mouse models show significant decline in treadmill performance. Mild, moderate, and severe FSHD-like mice were assessed for treadmill performance, as described in the methods. A) The moderately affected female mice (red line) were assayed prior to TMX injection (D0) and 3, 6, 10, 16, 23 and 29 days postinjection (DPI) and compared with age-matched female bi-transgenic mice (green line) and female *ACTA1-MCM* mice similarly injected with TMX (blue line). B) The moderately affected male mice (red line) were assayed prior to TMX injection (D0) and at 2, 6, 10, 14, 17, 21, 24 and 28 DPI and compared with aged-matched male bi-transgenic mice (green line) and male *ACTA1-MCM* mice similarly injected with TMX (blue line). C) Severely affected female mice (red line) were assayed prior to TMX injection (D0) and 2, 5, and 7, DPI (# D8, mice were too affected to be assayed) and compared with female bi-transgenic mice (green line) and *ACTA1-MCM* mice similarly injected with TMX (blue line). D) Severely affected male mice (red line) were assayed prior to TMX injection (D0) and 2, 5, 7, and 8 DPI and compared with age-matched male bi-transgenic mice (green line) and *ACTA1-MCM* mice similarly injected with TMX (blue line). Significance = * p<0.05, ** p<0.01, ***p<.001, calculated between bi-transgenic +TMX and bi-transgenic no TMX (green) or *ACTA1-MCM* +TMX (blue).

Treadmill assays indicated that muscle use and/or function was impaired by induction of *DUX4-fl* expression in the moderate and severe mouse models. *Ex vivo* muscle contractile studies using the isolated extensor digitorum longus (EDL) were then performed on these mice to assess changes in muscle strength [52, 53, 69]. Muscle forces from isometric contractions (twitch and tetanic) and tetanic force changes with increased frequencies of electric stimulation (force frequency) were measured and muscle stiffness was calculated from analysis of eccentric contractions (Figures 5 and S4). EDL muscles isolated from the mild model bi-transgenic mice (no TMX) were initially compared with age-matched *ACTA1-MCM* mice (control) treated with the appropriate dose of TMX for the respective group isolated at TMX D14 or TMX D10 from the first injection. Interestingly, despite having similar treadmill endurance profiles as controls (Figure 4), the female bi-transgenic mice (no TMX) consistently showed significantly lower maximum absolute force and specific force measurements (maximum force normalized by cross sectional area: CSA) compared with controls (Figure 5 and S4). Male bi-transgenic mice, however, were less affected and only showed a significant strength difference from controls with respect to maximum twitch force and maximum force frequency (Figure S4D-F), and no significant change in specific force measurements. However, both male and female bi-transgenic mild models were significantly less responsive to mid-level and high stimulation frequencies (65-180 Hz) compared with controls (Figure 5C and S4). We conclude that chronic, low, mosaic DUX4-FL expression in even a few myofibers reproducibly produced a measurable decrease in isometric contractile strength of muscle that does not appear to affect treadmill fitness. Thus, this data suggests that these mild bi-transgenic (no TMX) mice may present a model of pre-symptomatic FSHD or an early symptomatic mild FSHD.

**Figure 5:**
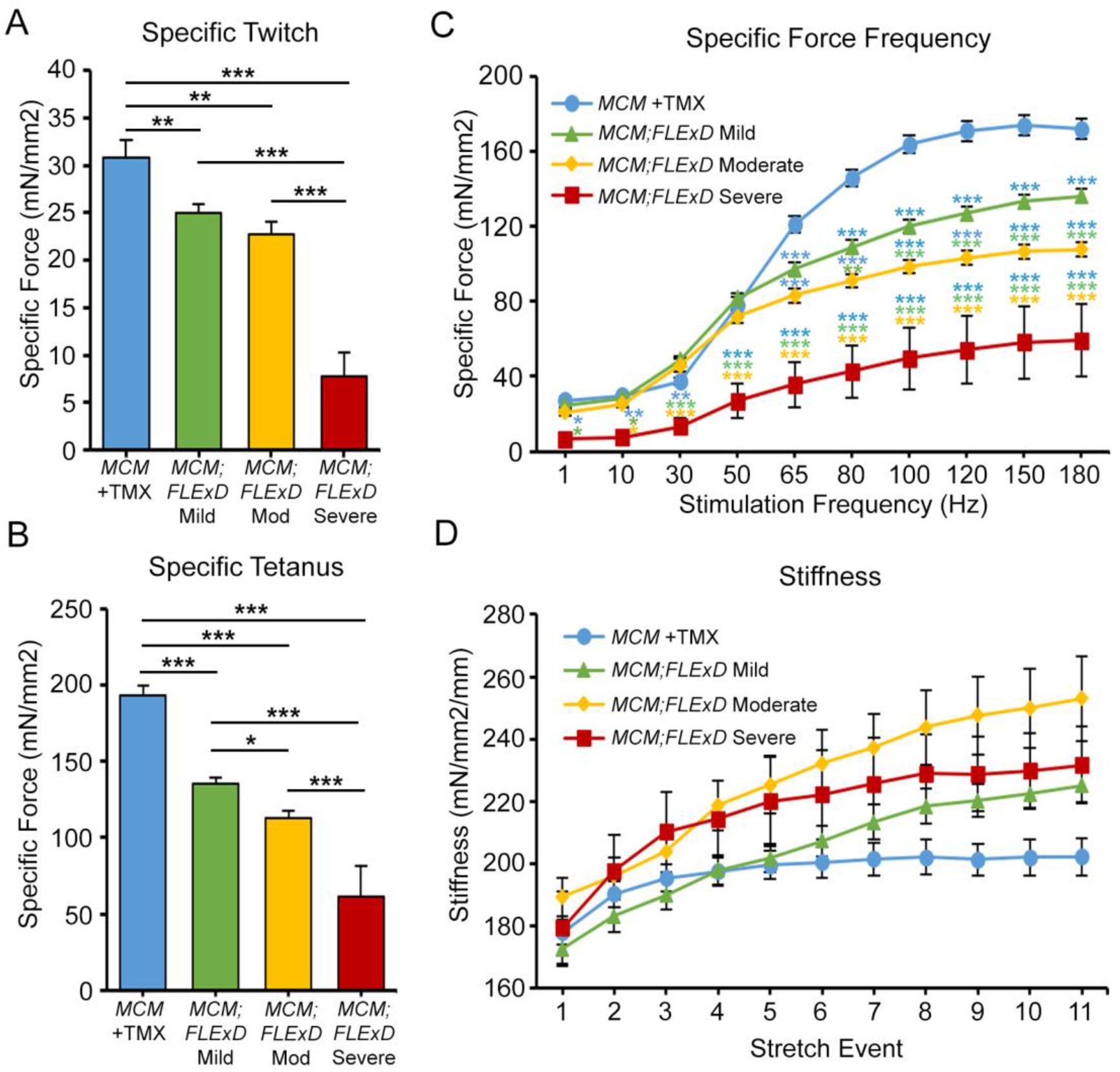
*Ex vivo* analysis of muscle function shows DUX4-induced muscle weakness. EDL muscles from female *ACTA1-MCM/+* with TMX (MCM, blue, n=9), bi-transgenic *ACTA1-MCM;FLExD* without TMX (*MCM;FLExD* Mild model, green, n=9), bi-transgenic *ACTA1-MCM;FLExD* with moderate TMX (*MCM;FLExD* Mod, yellow, n=6) and bi-transgenic *ACTA1-MCM;FLExD* with severe TMX (*MCM;FLExD* Severe, red, n=3) were assayed for maximum twitch, maximum tetanus, and force frequency (Figure S4), and then normalized to muscle cross sectional area to provide A) specific twitch, B) specific tetanus, and E) specific force frequency at MD14 and SD10. Significance = * p<0.05, ** p<0.01, ***p<.001; For panel C, blue asterisks indicate significance compared with *MCM* +TMX control mice and green asterisks indicate significance compared with mild bi-transgenic mice, yellow asterisks indicate significance compared with moderate model mice.

We similarly assayed the moderate and severe bi-transgenic mouse models for *ex vivo* muscle function, both of which showed significant differences in treadmill fitness compared with the *ACTA1-MCM* controls and the mild model mice upon TMX injection (Figure 4D and E). For the moderate model analysis, female mice were run on the treadmill until exhaustion, as above, prior to TMX injection, and then run twice per week until sacrificed and assayed at MD14, the peak of DUX4-FL protein expression (Figure 2D and I) and impaired treadmill running (Figure 4D) for the moderate model. The isolated EDL muscles from the moderate model mice were significantly weaker and stiffer compared with the *ACTA1-MCM* controls (Figure 5 and S4); however, these muscles only showed small, but significant, decreases in specific force measurements and stiffness when compared with the mild bi-transgenic mice. The DUX4-dependent impairment of muscle function was much more striking in the severe bi-transgenic model mice. Male and female mice were run on the treadmill until exhaustion, as before, prior to TMX injection and again at SD2, SD5, and SD7 (female) or SD8 (male), then sacrificed at SD10, the point of maximal fitness decline (Figure 4B and D) for the severe models. The EDL muscles from both female (Figure 5) and male (Figure S5) severe model mice showed ~3-fold decreases in all specific force measurements, and muscles were significantly stiffer after eccentric stretches when compared with *ACTA1-MCM* controls and the bi-transgenic mild model mice. In addition, the severe model mice were virtually non-responsive to low frequency stimuli (<30Hz) and were severely impaired across the force frequency assay range (Figure 5C and S5F). When compared with the moderate model mice, muscles from the severe model were again significantly weaker. Overall, we conclude that murine skeletal muscles can tolerate a very low level of DUX4-FL expression without significant impairment of function; however, increases in DUX4-FL expression eventually lead to muscle weakness and loss of function in a dose-dependent manner.

### Global mRNA expression analysis for moderate and severe FSHD-like mice

To begin to identify the mechanisms and pathways disrupted by DUX4-fl expression in murine skeletal muscle, RNA-seq analysis was performed on gastrocnemius muscles isolated from control, moderate (MD9), and severe (SD9) model mice, and analyzed for mRNA expression levels (Table S1). Genes with significant differential expression (>2-fold) from the *ACTA1-MCM* controls were identified (Table S2). The *ACTA1-MCM/+* and *FLExD/+* single transgenic mice only showed 3 genes differentially expressed (2 upregulated and 1 downregulated) between the mice (Figure 6A and Table S2), which is consistent with our previous qRT-PCR studies showing no significant differences in expression of *DUX4-fl* or several prominent DUX4-FL targets between these mice [41]. In contrast, the transcriptomes for both the moderate and severe models were significantly different from the *ACTA1-MCM* controls when analyzing genes significantly misregulated >2-fold (Figure 6B, C, and Table S2). The moderate model showed 1,479 genes differentially upregulated and 760 genes differentially downregulated, while the severe model showed 2,860 genes differentially upregulated and 1,882 genes differentially downregulated compared with *ACTA1-MCM* controls. This included known murine DUX4 target genes previously identified from C2C12 cells (Figure 6D-F) [23]. However, when these DUX4-induced genes (>2-fold) from murine skeletal muscle were compared with differential gene expression profiles from human FSHD patient muscle biopsies [24] there was very little overlap, with only 127 upregulated genes and 10 downregulated genes being the same (Table S3). This is not surprising since human DUX4 expressed in cultured mouse cells induces a gene set that only partially overlaps with DUX4 expression in cultured human cells [70, 71], and many functional DUX4 binding sites in the human genome are within poorly conserved retroelements [72].

**Figure 6:**
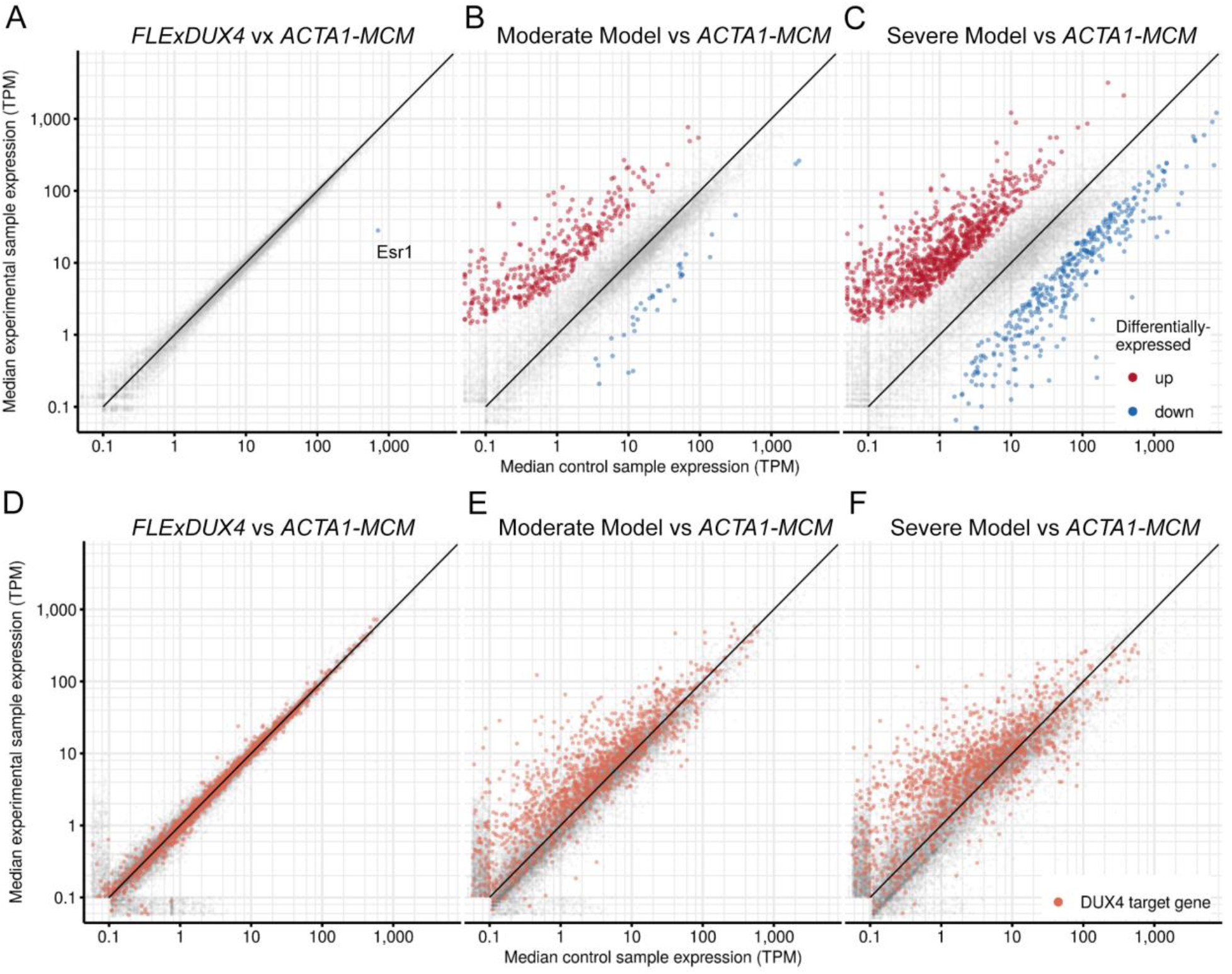
Comparison of gene expression in muscle from FSHD-like models compared with controls. (A-C) Scatter plots showing differentially expressed genes in RNA-seq data derived from tibialis anterior muscles isolated from A) *FLExDUX4/+* mice, B) bi-transgenic moderate model mice, and C) bi-transgenic severe model mice (y-axis), each compared with ACTA1-MCM control mice (x-axis). Significantly up-regulated genes are indicated in red, down-regulated genes are indicated in blue. (D-F) Above scatter plots highlighting the DUX4-induced genes similarly induced by DUX4 in C2C12 myoblasts [23].

Despite the low overlap in specific misregulated genes between muscles from FSHD patients and FSHD-like mice, there is some conservation in DUX4-activated pathways in human and mouse [23], and the general DUX4-induced myopathic phenotype appears to be conserved across several species [37, 73–76]. DUX4 is pro-apoptotic and its expression is detrimental to muscle development and differentiation [19, 34, 36–38, 73, 75, 77–80]. Expression of DUX4 stimulates genes that modulate the immune response [30], and FSHD patient biopsies show expression of immune genes, features of inflammation visualized by MRI, and immune cell infiltration [24, 81–84]. Therefore, we performed a gene ontology (GO) enrichment analysis on the differentially expressed genes from the FSHD-like moderate and severe mouse models focusing on pathways consistent with FSHD (Superterms: Apoptosis, Cell cycle, Immune, and Muscle; Figure 7, Tables 2, S4, and S5). A similar GO analysis using RNA-seq performed on C2C12 cells with inducible DUX4-fl expression [23] is shown for comparison and reveals differences between mouse muscle expressing human DUX4-fl and cultured murine cells expressing human DUX4-fl.

**Figure 7:**
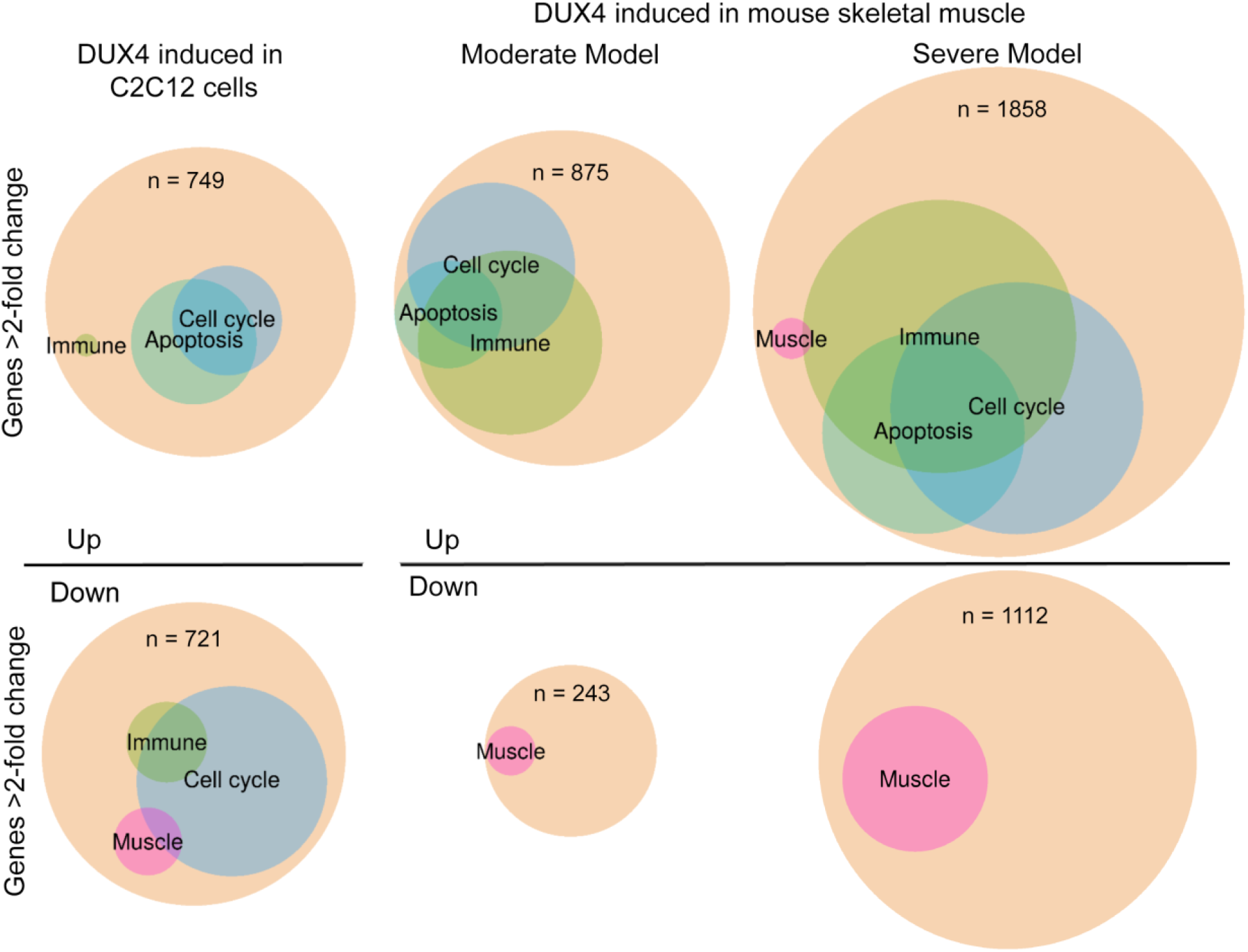
GO enrichment analysis of differentially expressed genes in FSHD-like model mice. GO enrichment analysis for DUX4-induced genes identified in C2C12 cells (left) [23], moderate FSHD-like model mice (middle) and severe FSHD-like model mice (right). The size of the beige-colored circles indicates the total number of differentially expressed genes (>2-fold), colored circles indicate the proportion of genes from significantly enriched GO-terms, that are in turn offspring of GO superterms (see methods and Table S4) relating to *mus*. Upregulated genes are represented in the top half and downregulated genes are represented in the lower half. The number of genes in each grouping is indicated (n).

DUX4-FL expression has been shown to inhibit nonsense-mediated decay, resulting in accumulation of mRNAs that would be typical substrates for NMD [33]. Therefore, we analyzed the RNA-seq data for aberrant alternative splicing events resulting in skipped exons (SE) or retained introns (RI) using MISO [56]. MISO identifies alternative splicing events based on predefined splicing events, taking advantage of existing knowledge and therefore is more accurate in characterizing alternative splicing events in RNA-seq data. The MISO analysis of the RNA-seq data revealed that the moderate and severe disease models had significant increases in SE and RI events and that the severe model had significantly more SE and RI events than the moderate model (Tables 3 and S6). This data supports that increased DUX4-fl expression and increased disease severity is correlated with increased SE and RI, which suggests a decrease in RNA quality control.

### Histological analysis of FSHD-like model mice

The three severity levels of myopathic mouse phenotypes (Table 1) were analyzed for DUX4-dependent histopathology. The initial analysis of the mouse models showed different, muscle-specific levels of transgene recombination in all three bi-transgenic severity models (Figure 1). Therefore, cryosections for histological analysis were generated from multiple muscles, including TA, gastrocnemius, soleus, quadriceps, and heart, for all three models. These muscle sections were then analyzed by hematoxylin and eosin (H&E) staining to assess fiber morphology, TUNEL assay to assess apoptosis, embryonic myosin heavy chain (eMyHC/Myh3) IF to assess muscle fiber regeneration, picrosirius red (SR) staining to assess fibrosis, and CD11b and Ly6G IF to assess immune cell infiltration. The previously described time-courses of treadmill exhaustion running were performed using female mice for the moderate model, with mice sacrificed for histology at MD3, MD6, MD14, and MD28 (Figure 8), and using male mice for the severe model, with mice sacrificed at SD3, SD6, and SD9 (Figure 9). Sex and age-matched *ACTA1-MCM* mice injected with appropriate levels of TMX for the model were used as controls and the mild bi-transgenic (No TMX) model mice were assayed for comparison (D0).

**Figure 8:**
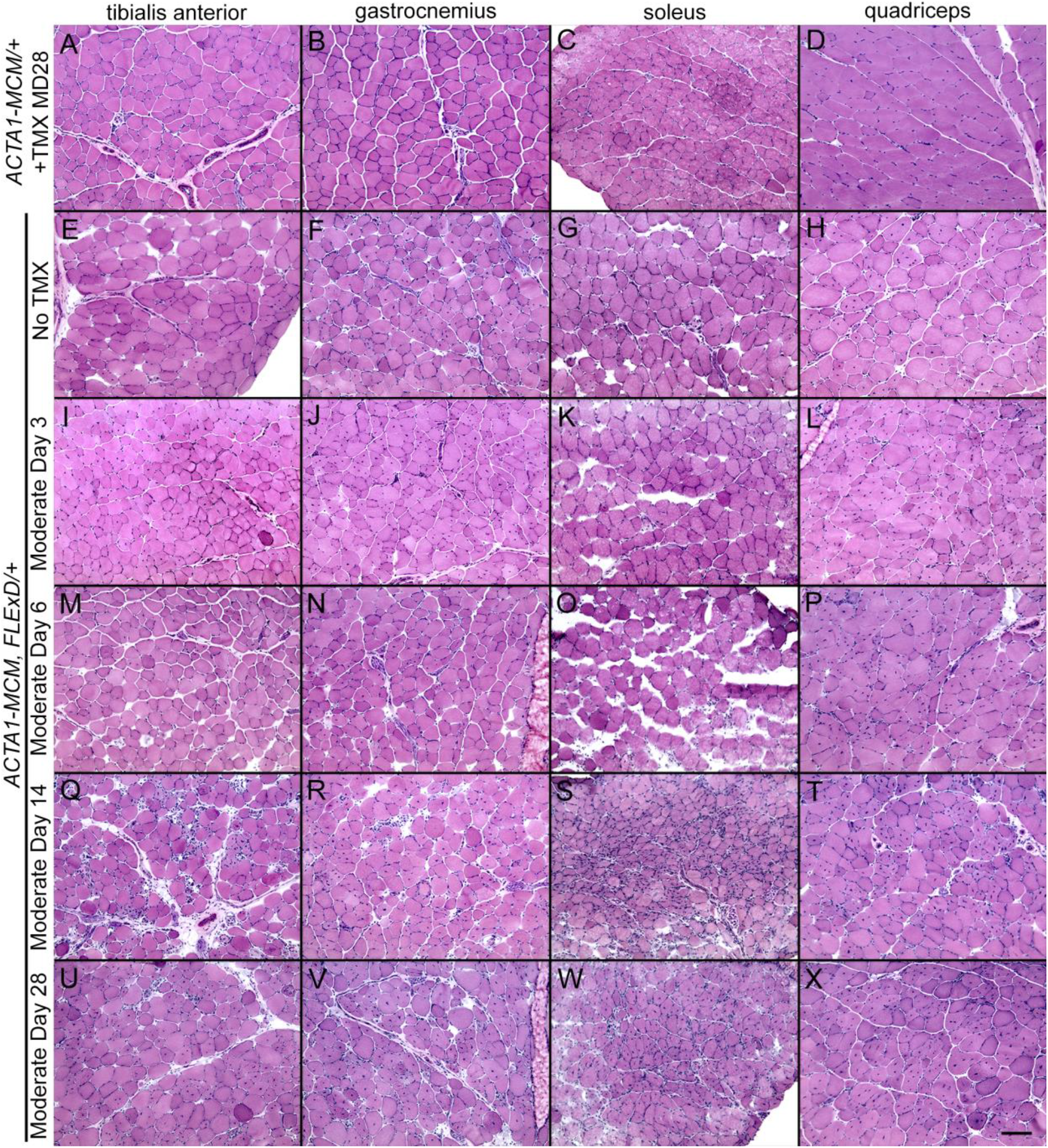
Skeletal muscles from the mild and moderately affected FSHD-like mouse models show signs of histopathology. Representative cryosections of tibialis anterior, gastrocnemius, soleus, and quadriceps of indicated transgenic animals were analyzed with H&E staining. Female (A-D) *ACTA1-MCM/+* mice treated with 1X 5mg/kg TMX, (E-H) mild model bi-transgenic mice without TMX, and (I-X) moderate model bi-transgenic mice treated 1X with 5mg/kg TMX. The TMX-injected *ACTA1-MCM* control mice were assayed at MD28. The TMX-injected bi-transgenic mice were assayed at (I-L) MD3, (M-P) MD6, (Q-T) MD14, and (U-X) MD 28. All mice had performed the treadmill exhaustion analysis, as described, 2X per week starting the week prior to TMX injection. Scale bar = 100 μm

**Figure 9:**
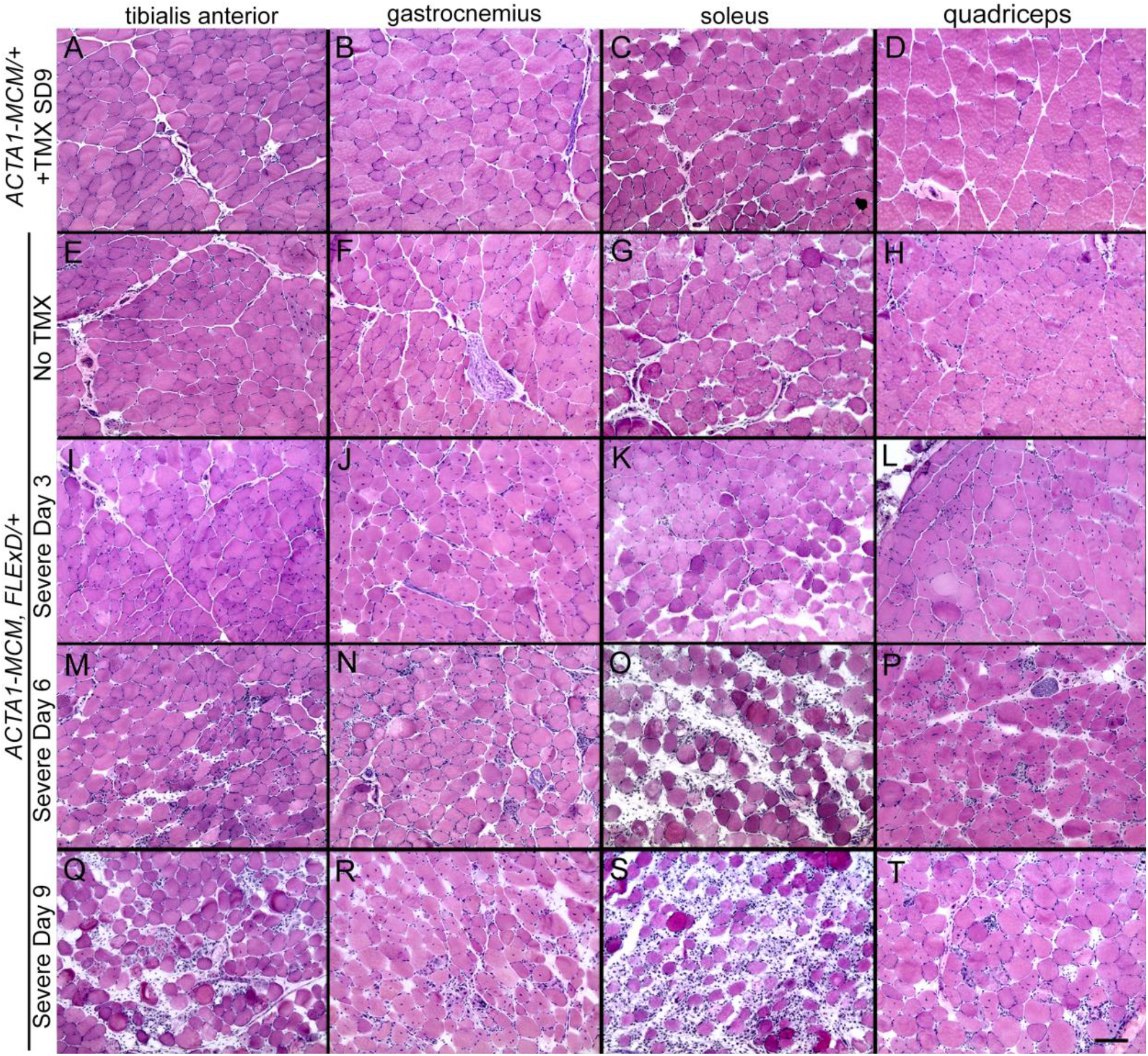
Skeletal muscle from the severely affected FSHD-like mice show signs of a severe myopathy. Representative cryosections of tibialis anterior, gastrocnemius, soleus, and quadriceps from indicated transgenic animals were analyzed with H&E staining. A-D) *ACTA1-MCM*/+ mice treated with 2X 10mg/kg TMX, (E-H) mild model bi-transgenic mice without TMX, and (I-T) severe model bi-transgenic mice treated 2X with 10mg/kg TMX. The TMX-injected controls were assayed at SD9. The TMX-injected bi-transgenic mice were assayed at (I-L) SD3, (M-P) SD6, and (Q-T) SD9. All mice had performed the treadmill exhaustion analysis, as described, 1 week prior to TMX injection, then 2, 5, and 7 DPI. Scale bar = 100 μm

Analysis of the H&E histology indicated that for the mild model muscles (Figure 8E-H and Figure 9E-H), very low mosaic DUX4-fl expression leads to minor changes in histology, the most notable being increased centralized nuclei compared with *ACTA1-MCM* control muscles (Figure 10). Interestingly, there are anatomical muscle-specific effects that correlate with the level of transgene recombination. The soleus muscles, which have the lowest levels of leaky transgene recombination in muscle assayed for the mild model, have ~3% myofibers with central nuclei, while the TA muscles, which have a higher recombination rate and thus more DUX4-FL-expressing nuclei (Figure 1B), have ~10% myofibers with central nuclei (Figure 10), suggesting that DUX4-FL expression is driving the formation of fibers with central nuclei.

**Figure 10:**
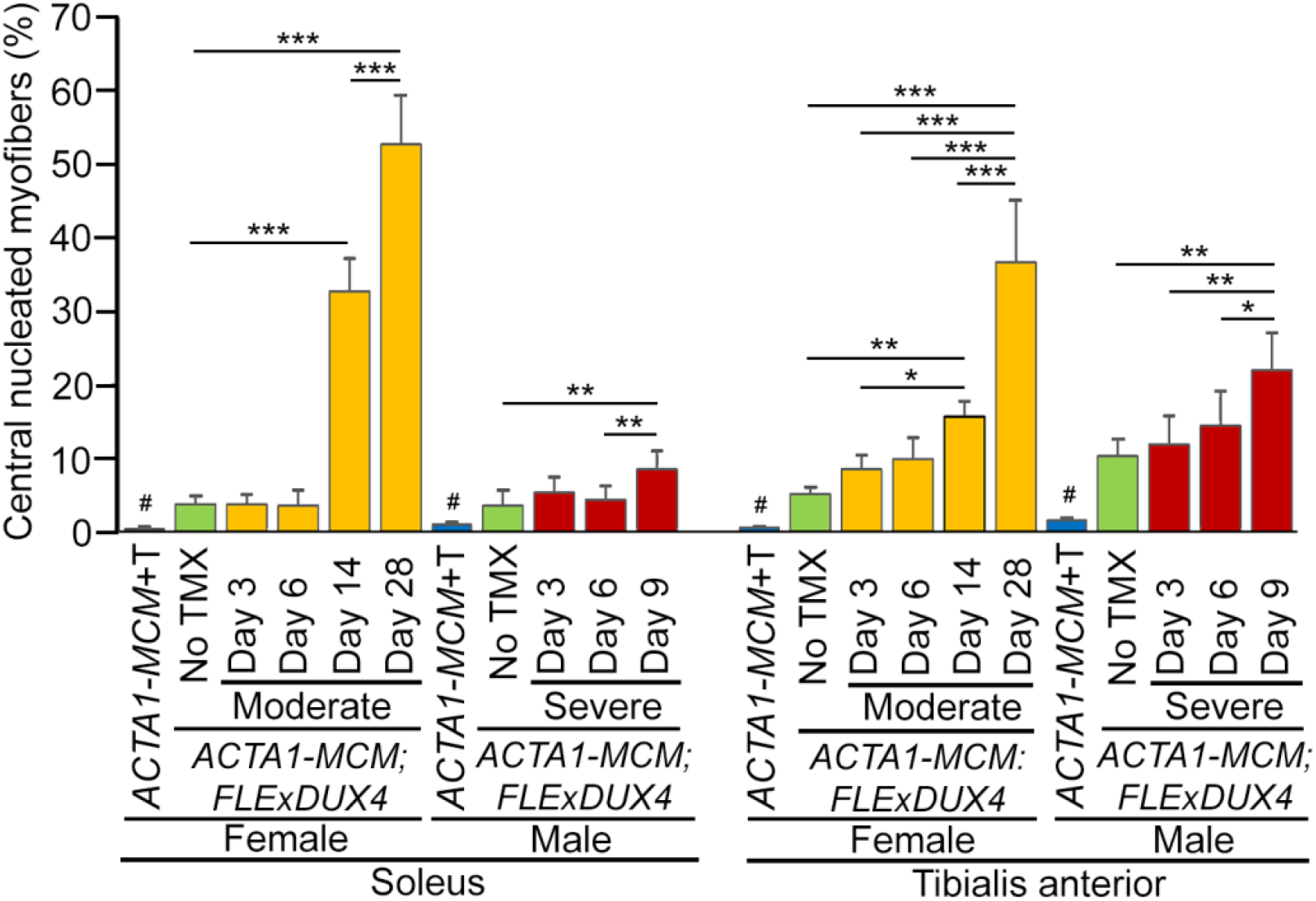
Skeletal muscles from FSHD-like mouse models develop centralized nuclei. H&E stained histological cross sections from soleus and tibialis anterior muscles were dissected from *ACTA1-MCM* control (blue), and mild (green), moderate (yellow), and severe (red) FSHD-like model mice that had undergone treadmill exhaustion analysis 2X per week (see Figures 7 and 8). Muscles were assayed from bi-transgenic mice without TMX injection (No TMX) for the mild model, from female bi-transgenic mice at MD3, MD6, MD14, and MD28 for the moderate model, and from male bi-transgenic mice at SD3, SD6, and SD9 for the severe model mice. The *ACTA1-MCM* +TMX controls were assayed at MD28 and SD9, as appropriate. Data is plotted as percent of myofibers with centralized nuclei. Significance was calculated by XXX, *= p<.05, **= p<.01, ***= p<.001, # = p<.001 for *ACTA1-MCM* compared with all other samples in the group.

In contrast to the mild model, the histology from the moderate (Figure 8I-X) and severe (Figure 9I-T) model mice showed greater effects of DUX4-fl expression, including variability in skeletal muscle fiber size, round and triangular fiber shapes, significant increases in centralized nuclei, an influx of mononuclear cells, and ultimately an apparent decrease in structural integrity of the muscle. This histopathology accumulated over time and correlated with loss of muscle function. For example, in the moderate model histology, the few mononuclear cells that have infiltrated by MD6 dramatically increase by MD14. Similarly, the percentage of fibers with centralized nuclei jumps significantly between MD6 and MD14 and MD28 in both the soleus and TA (Figure 10). Importantly, as seen in the mild model, these models showed anatomical muscle-specific differences in histopathogenic features, correlating with muscle-specific levels of transgene recombination (Figure 1B), consistent with DUX4-FL expression leading to an atrophic phenotype [34]. For example, the soleus muscle, which shows the highest level of TMX-responsive transgene recombination, also appears to have the greatest degree of histopathology for each model, including mononuclear cell infiltration, disrupted muscle integrity (Figure 8K, O, S and Figure 9O, S), and fibers with centralized nuclei (Figure 10). The quadriceps showed the lowest level of transgene recombination and similarly showed the least amount of histopathology for the analyzed skeletal muscles. The heart, which showed no transgene recombination or expression, did not show any pathology even in the severe model (Figure S6). Overall, the skeletal muscle histopathology in these models was progressive, appearing with peak DUX4-FL expression levels at MD14 in the moderate model while appearing earlier in the severe model (SD6), and peaking with DUX4-FL levels at SD9. This further supports that the extent of pathology in these mouse models correlates with the level of transgene recombination (Figure 1B) and DUX4-fl expression (Figure 2 and 3).

The centrally positioned nuclei in myofibers are a common feature of many myopathies and are considered a sign of repair and regeneration of the myofiber [85]. Interestingly, all three severity models of FSHD-like mice show increased centralized nuclei, especially following DUX4-FL induction (Figure 10). Therefore, we analyzed muscles from these models for expression of eMyHC protein (Figure 11), a marker for newly regenerating myofibers and dystrophic muscle [86], and expression of *Myostatin (Mstn)/Gdf-8* (Figure S7), a negative regulator of muscle growth and remodeling in adult muscle [87]. Adult mouse skeletal muscles have very few myofibers that express the eMyHC isoform under normal healthy conditions [88]. However, regenerating myofibers re-express many developmental isoforms of muscle proteins, including eMyHC, which can be detected within 3 days of injury and whose expression can persist for up to 3 weeks [89–91]. Therefore, a time course study of eMyHC expression after DUX4-fl induction was performed on TA muscles (Figure 11). As expected, the *ACTA1-MCM* controls showed no detectable expression of eMyHC in response to TMX (Figure 11A-C). Interestingly, the mild model bi-transgenic mice, which have low mosaic expression of DUX4-FL, similarly showed no detectable expression of eMyHC (Figure 11K, L, S, T), indicating that myofibers with centralized nuclei in the mild model are remnants of an old regeneration event. In contrast, both the moderate model (Figure 11E-J) and severe model (Figure 11M-R) showed high levels of eMyHC, peaking at MD14 and SD9, respectively, thus confirming that the spike of DUX4-FL expression at these moderate and severe levels activates the skeletal muscle regeneration program. In contrast to eMyHC levels, expression of *Mstn* mRNA decreases in TA muscles as DUX4 levels and severity of pathology increased (Figure S7). Together, these results show increased DUX4 expression leads to the muscle remodeling and regeneration.

**Figure 11:**
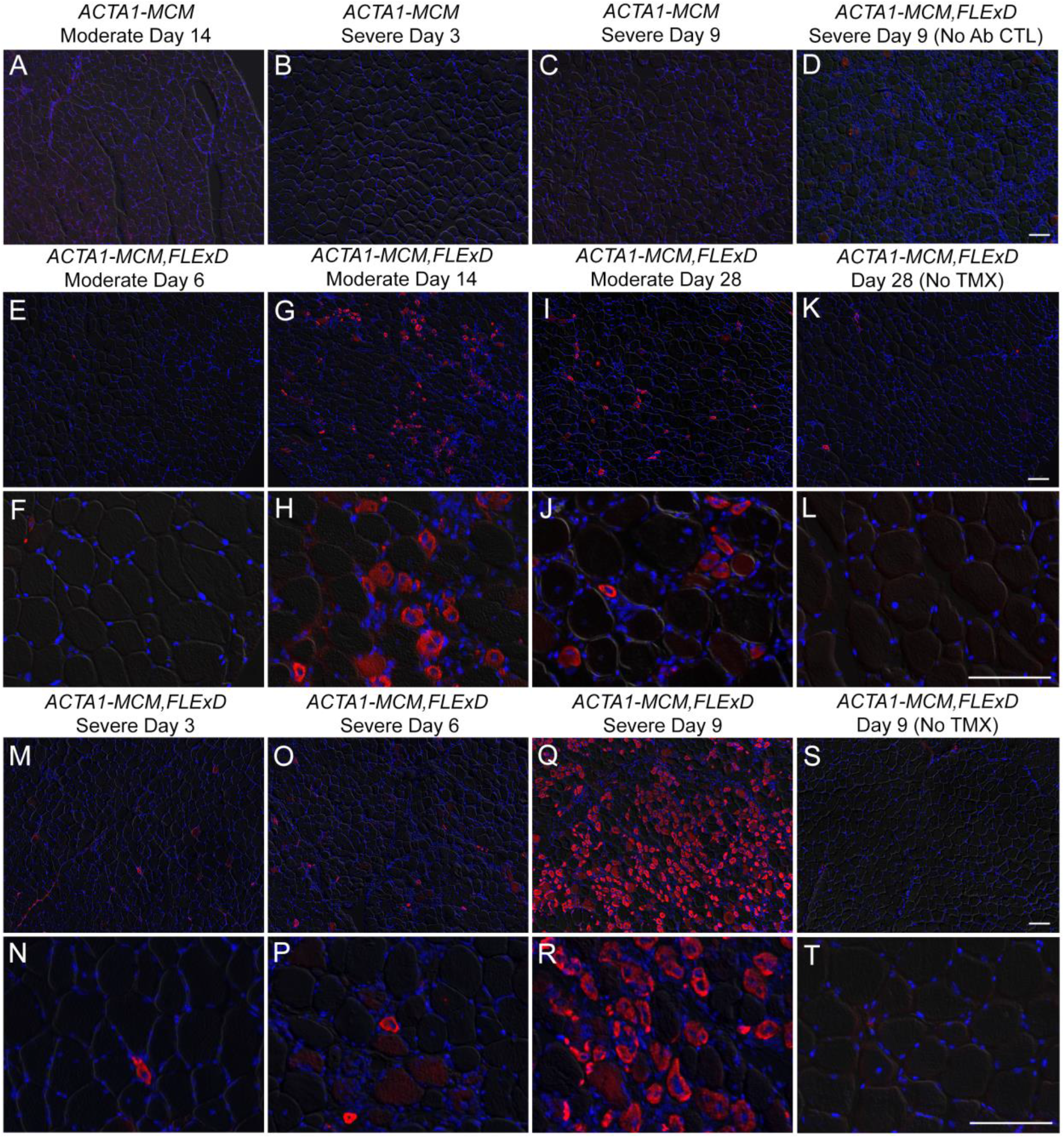
Induced DUX4-fl expression causes regeneration of muscle fibers marked with eMyHC expression. Histological cross sections from tibialis anterior muscles were dissected from mild, moderate, and severe FSHD-like model mice that had undergone treadmill exhaustion analysis 2X per week (see Figures 7 and 8). Age-matched female control mice were used for the moderate model analysis and male control mice for the severe model analysis. Muscles were assayed from *ACTA1-MCM*/+ TMX controls at (A) MD14, (B) SD3, and (C) SD9. Muscles were assayed from mild model bi-transgenic mice without TMX injection at (K, L,) 28 DPI and (S, T) 9 DPI, from female bi-transgenic mice at (E, F) MD6, (G, H) MD14, and (I, J) MD28 for the moderate model, and from male bi-transgenic mice at (M, N) SD3, (O, P) SD6, and (Q, R) SD9 for the severe model. Scale bars = 100 μm

The activation of muscle fiber regeneration after induction of DUX4-FL expression suggests that DUX4-FL is causing muscle damage. The GO analysis of differentially induced genes showed that muscles from the moderate and severe models are enriched for genes in apoptotic pathways (Figure 7, Table 2). As mentioned, DUX4-FL is a pro-apoptotic protein, its expression is highly toxic to muscle cells in culture [19, 36, 77, 92], and apoptosis is a feature of FSHD muscle [93]. To assess apoptosis in the FSHD-like models, TUNEL assays were performed on TA muscles across a DUX4-FL induction time-course in both the moderate and severe models (Figures 12 and 13, respectively). In the moderate model, TUNEL-positive nuclei appeared by MD6, were prevalent at MD14, and were nearly absent by MD28 (Figure 12A-C). Similarly, the severe model showed TUNEL positive nuclei by SD6, with a dramatic increase by SD9 (Figure 13A-C). Interestingly, muscles from the bi-transgenic mild model showed no indication of apoptosis and were similar to muscles from *ACTA1-MCM* controls (Figure13, compare panels D and L). Thus, the moderate and severe models show dose-dependent DUX4-FL-induced apoptosis following a similar time course as muscle weakness and decreased function, and correlating with the activation of muscle regeneration.

**Figure 12:**
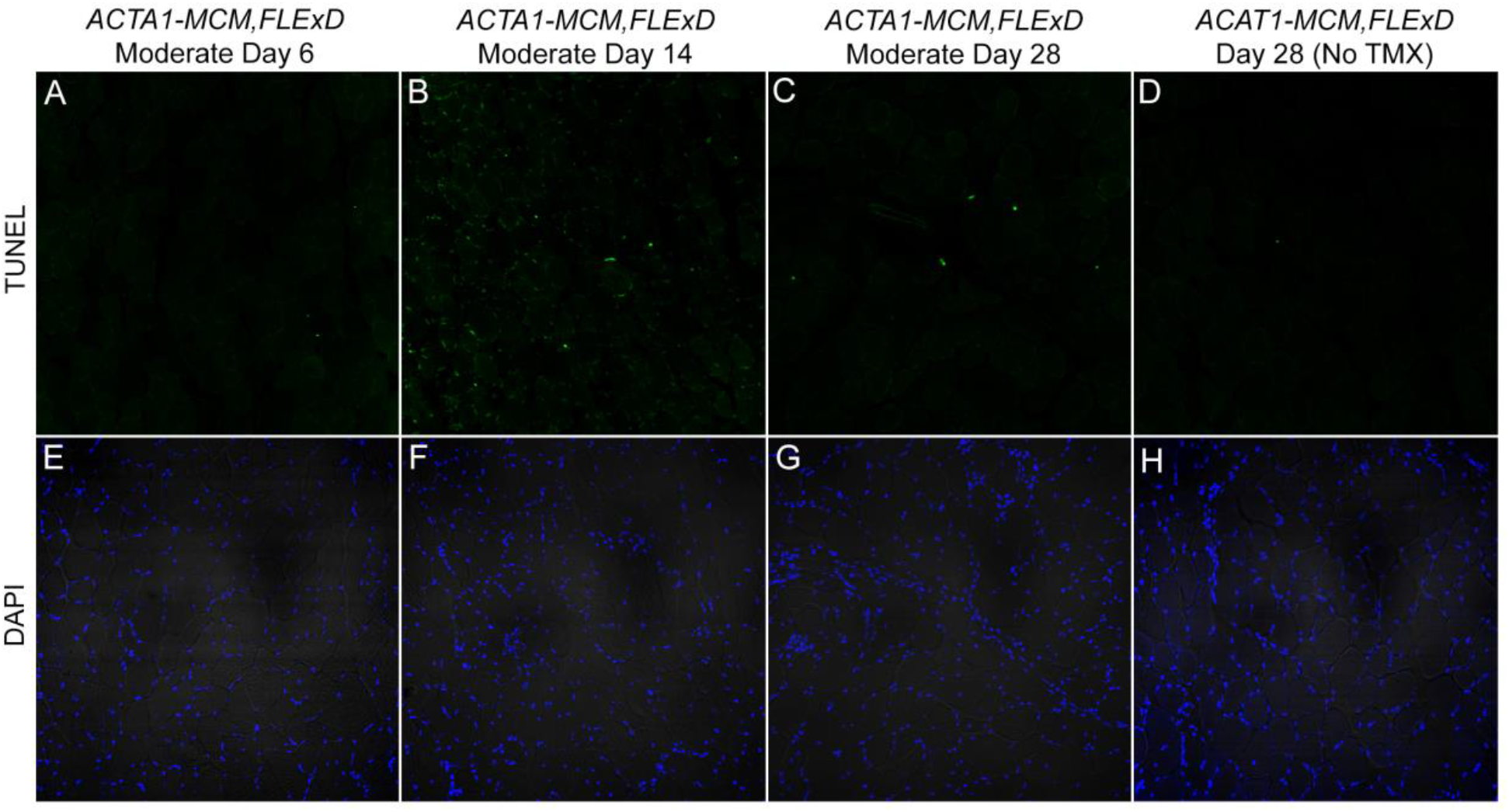
Muscles from the moderate FHSD-like model mice undergo apoptosis in response to induced DUX4-fl expression. Tibialis anterior muscles from female TMX-injected bi-transgenic moderate model mice that had undergone treadmill exhaustion analysis 2X per week (see Figure 8) were assayed at MD6 (A, E), MD14 (B, F), and MD28 (C, G) or at MD28 for matched non-injected bi-transgenic mice (D, H). Green signal in the TUNEL assay (A-D) indicates nuclei undergoing apoptosis, compared with DAPI staining (E-H) showing all nuclei in the same histological sections.

**Figure 13:**
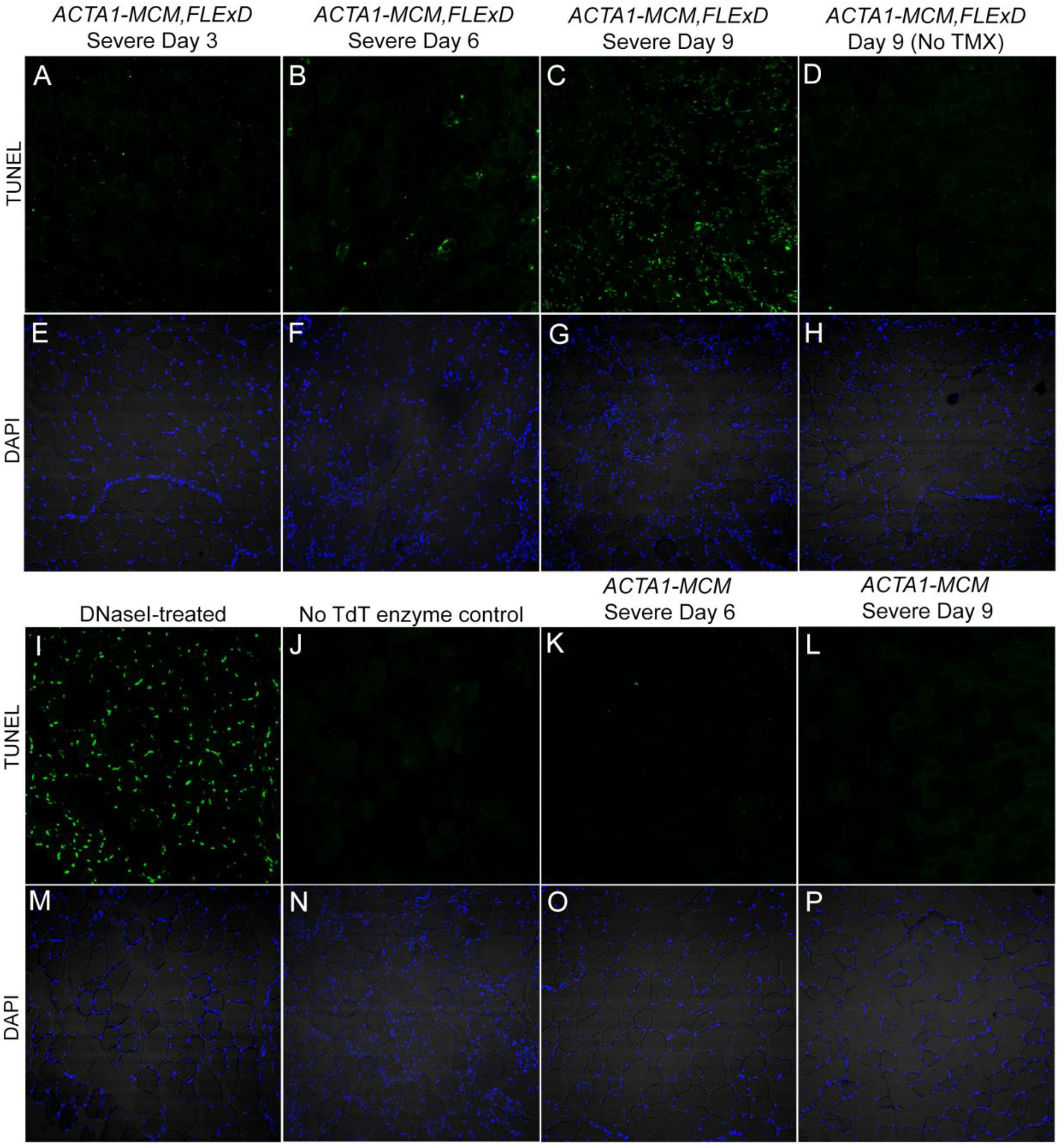
Muscles from the severe FSHD-like model mice start to undergo apoptosis within 6 days of induced DUX4-fl expression. Tibialis anterior muscles from male TMX-injected bi-transgenic severe model mice that had undergone treadmill exhaustion analysis (see Figure 9) were assayed at SD3 (A, E), SD6 (B, F), and SD9 (C, G) or at SD9 for matched noninjected bi-transgenic mice (D, H). Green signal in the TUNEL assay (A-D, I-L) indicates nuclei undergoing apoptosis, compared with DAPI staining (E-H, M-P) showing all nuclei in the same histological sections.

Patient biopsies show that immune cell infiltration and inflammation are features of FSHD muscle [24, 81–84, 94]. Similarly, the RNA-seq analysis of muscle from the moderate and severe FSHD-like mouse models indicated an enrichment for immune response genes (Figure 7), and the H&E staining showed mononuclear cell infiltrates after DUX4 induction and the appearance of histopathology (Figures 8 and 9). Therefore, the mononuclear cell infiltrates in gastrocnemius muscles from the moderate (Figure 14) and severe (Figure 15) models were assayed over the time courses of DUX4 induction. Muscle sections were immunostained for CD11b (*Itgam*, Integrin subunit alpha M), a member of the CD18 integrin family of leukocyte adhesion receptors involved in leukocyte migration [95], and Ly6g (lymphocyte antigen 6 complex, locus G), a marker for murine neutrophils [96, 97]. IF confirmed the presence of these pro-inflammatory cells in the affected muscles of the moderate model by MD14 of DUX4-fl induction (Figure 14J-L), which disappeared by MD28. In the severe model, immune cells appeared as early as six days (SD6) after *DUX4-fl* induction (Figure 15G-I) and by SD9, muscles were severely affected, showing large accumulations of immune cell infiltration (Figure 15J-L). These immune cell infiltrates were absent in muscles from the mild model and from *ACTA1-MCM/+* mice (Figure 14A-C and Figure 15A-C, respectively). Importantly, these immune phenotypes in the moderate and severe models are consistent with pathology seen in muscles from clinically affected FSHD patients, while the lack of immune cell infiltration in the mild model is consistent with the lack of overt pathology in pre-symptomatic or asymptomatic FSHD subjects.

**Figure 14:**
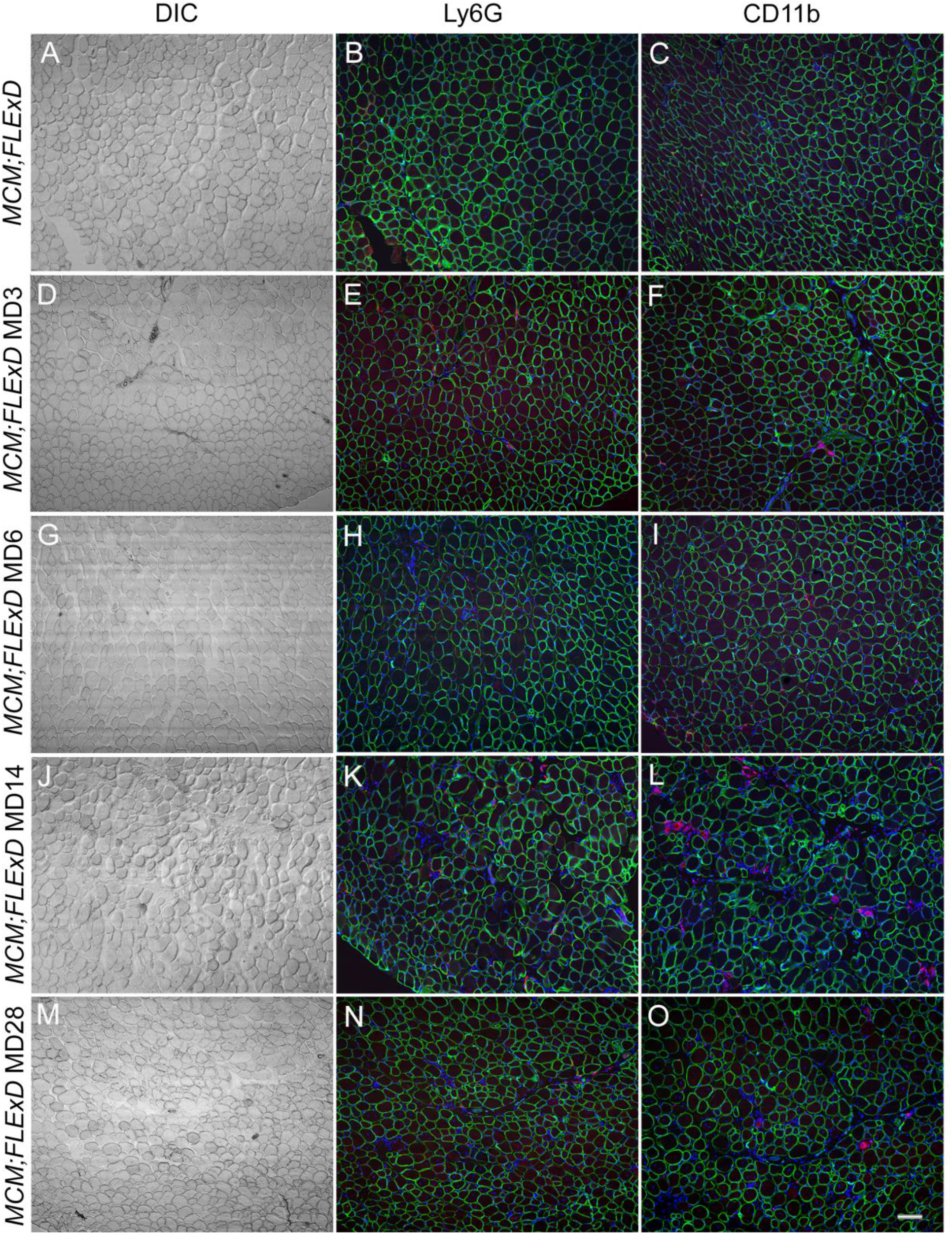
Induction of *DUX4-fl* leads to innate immune cell infiltration in skeletal muscles of moderate but not the mild model FSHD-like mice. Gastrocnemius muscle sections from AC) mild model bi-transgenic mice and moderate model mice at D-F) MD3, G-I) MD6, J-L) MD14, and M-O) MD28 were immunostained for Ly6G (red, middle column) or CD11b (red, right column) and dystrophin (green), to reveal immune cell infiltration upon TMX induction of DUX4-fl expression. DAPI (blue) shows the nuclei. Images at 10X magnification, scale bar = 100μm

**Figure 15:**
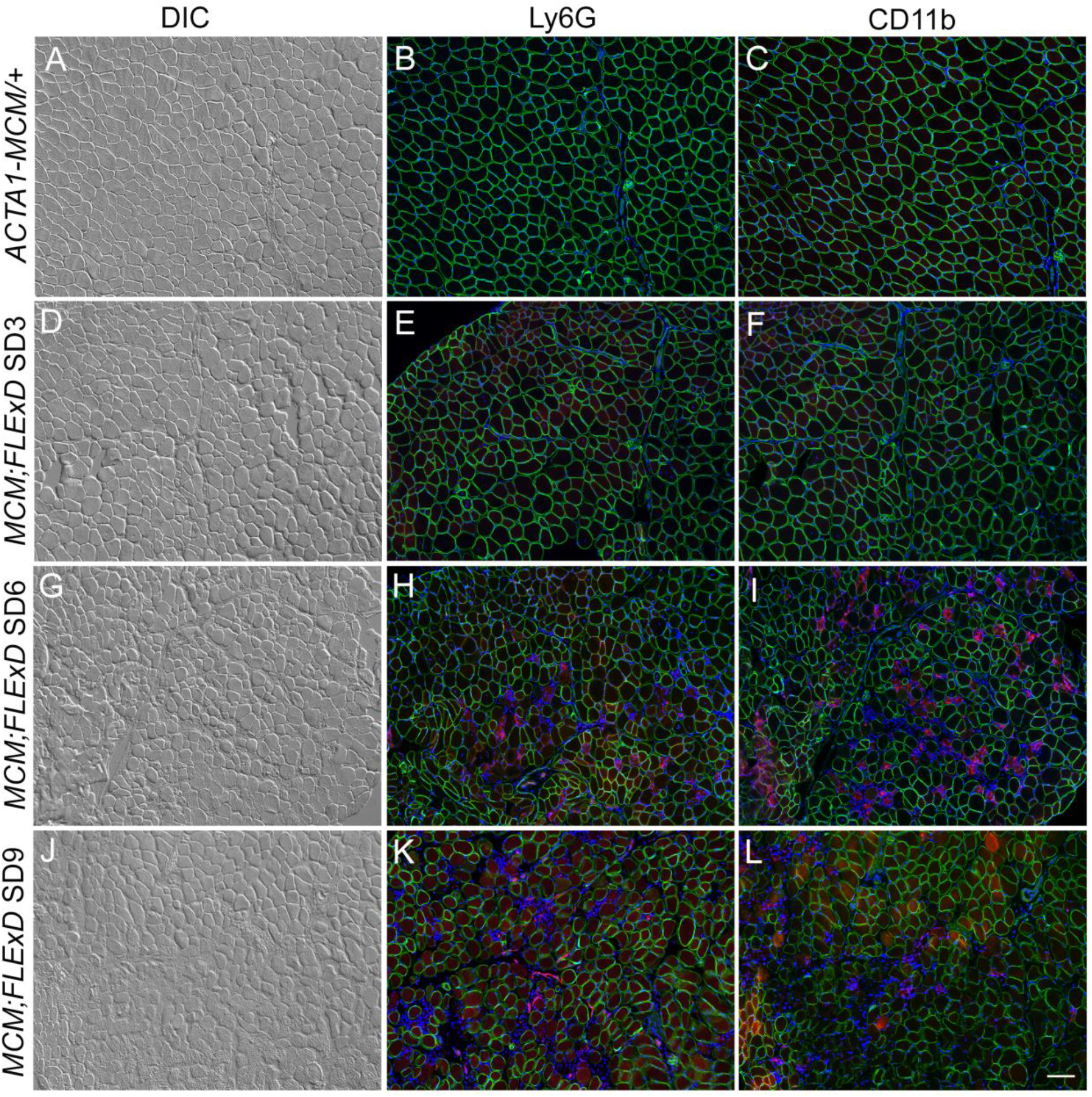
Induction of *DUX4-fl* leads to innate immune cell infiltration in skeletal muscles of severe model mice. Gastrocnemius muscle sections from control A-C) *ACTA1-MCM*, and severe model mice at D-F) SD3, G-I) SD6, and J-L) SD9 were immunostained for Ly6G (red, middle column) or CD11b (red, right column) and dystrophin (green) to reveal immune cell infiltration upon TMX induction of DUX4-fl expression. DAPI (blue) shows the nuclei. Images at 10X magnification, scale bar = 100μm

Muscle biopsies and MRI data indicate increased fibrosis and fatty replacement in FSHD patients [84, 94]. Muscle fibrosis, deposits of extracellular matrix between myofibers, is another feature of many muscular dystrophies, including FHSD, and leads to the loss of muscle architecture and decreased muscle function [28, 98–101]. Inflammatory cells, such as those present in the FSHD-like mouse models (Figures 14 and 15), are able to stimulate fibrotic activity during regeneration and promote fibrotic tissue remodeling [99, 102]. The extent of fibrosis is typically quantified using histological methods and staining for collagen [51, 103]. Therefore, cross sections of the TA muscles isolated from the above series of control, mild, moderate, and severe FSHD-like mice were assayed for the extent of fibrosis developing over time using SR staining (Figures 16, S8, and S9). The muscles from control *ACTA1-MCM/+* and mild bi-transgenic model had similar low levels (~2% fibrotic area) of fibrosis (Figure 16A, F, G, and Figure S8). The moderate bi-transgenic model had similar control levels of fibrosis at MD3 and MD6; however, this model showed a small but significant 50% increase in fibrosis by MD14 and MD28 (Figure 16B-E and Figure S8). The severe bi-transgenic model showed increased fibrosis by SD3 and a maximal 2.5-fold increase in fibrosis (5.5% fibrotic area) at SD9 (Figure 16 H-J, and Figure S8). The heart, which does not express detectable DUX4-fl in any of the models, showed no signs of fibrosis (Figure S9).

**Figure 16:**
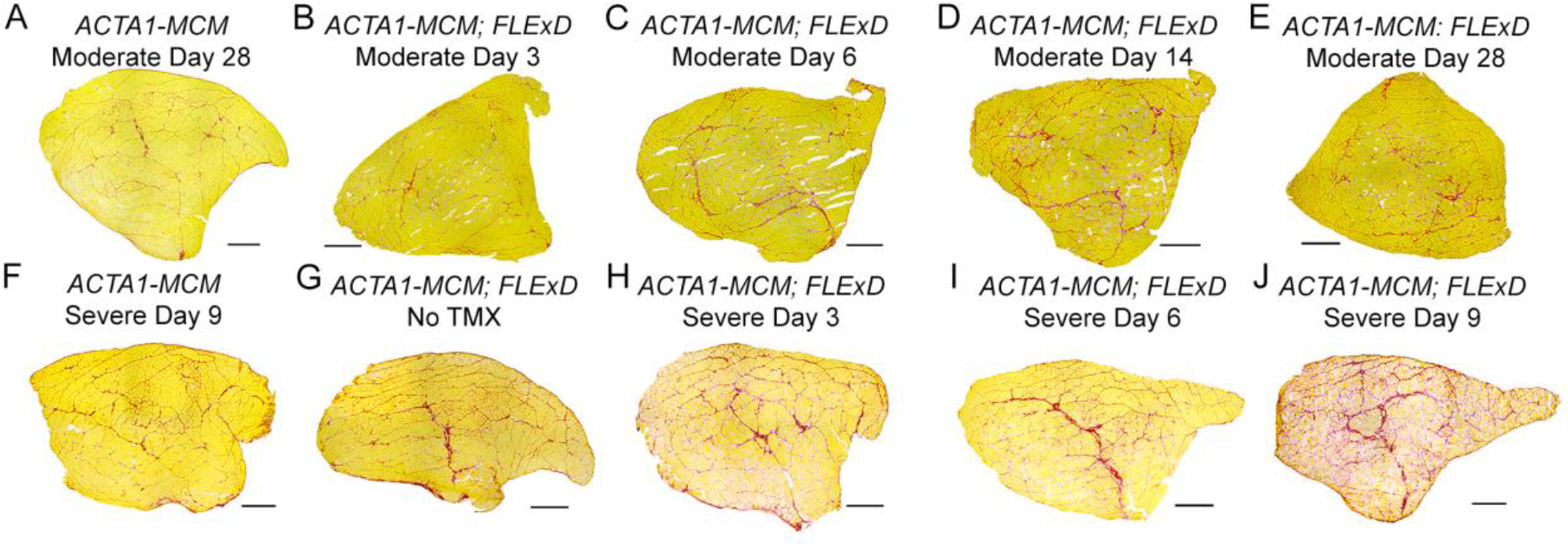
Increased DUX4-fl expression in the moderate and severe FSHD-like model mice leads to accumulation of fibrotic tissue in skeletal muscles. Histological cross sections from tibialis anterior muscles were dissected from mild, moderate, and severe FSHD-like model mice that had undergone treadmill exhaustion analysis 2X per week (see Figures 7 and 8). Age-matched female control mice were used for the moderate model analysis and male control mice for the severe model analysis. Muscles were assayed from *ACTA1-MCM*/+ TMX controls at MD28 (A) and SD9 (B), from bi-transgenic mice without TMX injection at MD28 (G), from female bi-transgenic mice at MD3 (B), MD6 (C), MD14 (D), and MD28 (E) for the moderate model, and from male bi-transgenic mice at SD3 (H), SD6 (I), and SD9 (J) for the severe model. Representative histology images are shown. Quantitative analysis of data shown in Figure S8. Scale bars = 500 μm

## Discussion

Modeling FSHD in transgenic mice has historically been very difficult despite the fact that FSHD is a gain-of-function disease seemingly amenable to transgenic *DUX4* overexpression [68, 74, 76]. While the human *DUX4* gene has a conserved developmental role with *Dux* family members found in other mammals, including the mouse, there is significant divergence at the DNA and protein sequence level as well as in the spectrum of species-specific target genes [23, 70, 104]. In addition, because DUX4-FL is highly cytotoxic for many somatic cells, leaky expression during development has been problematic, and surviving mice can be severely phenotypic and difficult to breed [75]. However, the *FLExD* mice we recently developed overcame many of these previous limitations [41]. Both male and female mice are fertile and easy to breed, male or female transgenic mice can be produced either as transgene heterozygotes or homozygotes, which can live more than 1.5 years, and, when mated with an inducible Cre line of mice, bi-transgenic mice allow for investigator-controlled expression of DUX4-fl mRNA and protein resulting in FSHD-like muscle pathology [41]. Here we report the characterization of a series of phenotypic FSHD-like mouse models varying in severity from mild to severe, generated using the *FLExD* conditional *DUX4-fl* transgenic mouse line crossed with *ACTA1-MCM* TMX-inducible mice. We demonstrate that these bi-transgenic mouse models, with and without TMX induction, recapitulate many aspects of FSHD pathophysiology with variable severity, thus providing suitable models for therapeutic interventions targeting DUX4-fl mRNA, protein, and certain downstream pathways, with several key caveats. In particular, it is important for those working on these models to keep in mind the described anatomical muscle-specific and sex-specific differences in pathology and disease progression.

### Mild FSHD-like mouse model

The *ACTA1-MCM;FLExD* bi-transgenic mouse, in the absence of any TMX induction, has mosaic expression of DUX4-fl mRNA and protein and provides an excellent model of mild, pre-symptomatic FSHD. This is because the TMX-inducible Cre fusion protein produced in skeletal muscles by the *ACTA1-MCM* transgenic line exhibits leakiness into nuclei, resulting in Cre activity in the absence of TMX in a fraction of nuclei. Thus, in the *ACTA1-MCM;FLExD* bi-transgenic mice, this results in recombination of the DUX4 transgene and low mosaic expression of DUX4-FL protein in skeletal muscles throughout their lifetime. Importantly, although this leaky recombination is specific to skeletal muscle, it is not uniform among skeletal muscles, with different anatomical muscles exhibiting different, but consistent, levels of recombination and thus DUX4-fl expression (Figure 1B, e.g. soleus is low and quadriceps is high). Thus, the situation in this mild model is similar to that in pre-symptomatic or asymptomatic FSHD patients. These mice live with a chronic, low-level, mosaic expression of DUX4-FL in a fraction (<10%) of muscle fibers, yet phenotypically the mice appear healthy and behave normally, with no changes in overall fitness or lifespan.

Although these mild mice appear outwardly healthy, the low-level recombination produces several assayable phenotypes. The mild phenotype manifests as increased expression of DUX4-FL target genes (Figure 3), decreased expression of *Mstn* (Figure S7), 5-10% of myofibers with centralized nuclei at 10-12 weeks of age (Figures 7 and 8), and 40% decreased capacity for muscle force generation in female (but not male) mice, assayed *ex vivo* (Figures 5, S4 and S5). There are also small but significant differences between males and females with respect to histology and centralized nuclei that need to be taken in to account. Overall, there is some low-level muscle pathology and new fiber regeneration, however, the mice do not exhibit increased apoptosis, immune cell infiltration, or increased fibrosis. The model is readily scalable, highly reproducible, and, considering that these mice live >1.5yrs (female n=43, male n=20), provides a DUX4-fl expression model that is amenable to longevity studies for efficacy of putative DUX4-targeted therapeutics. Thus, while the mild model is likely the easiest to work with and imposes no time limits on treatments, investigators should be careful in choosing which sex to use (or use a significant number of both sexes and analyze them separately), which age to use, and which muscles to assay or treat.

### Moderate FSHD-like mouse model

Similar model-specific effects need to be taken into account when injecting TMX to generate the moderate and severe models, but these are even more pronounced. Importantly, while the moderate model showed consistently different levels of TMX-induced recombination between anatomical muscles (Figure 1B), these were different from the leaky recombination in the mild model and more likely a reflection of the difference in TMX accessibility to various muscles. The moderate model mice also showed significant differences between male and females with respect to weight (Figure S3), treadmill profiles (Figure 4), and muscle physiology (Figures 5 and S4), with females being more severely affected by all metrics. With respect to appearance and progression of pathology, the mice show further reduction in *Mstn* expression, and, as opposed to the mild model, the moderate model mice showDUX4-induced apoptosis (Figure 12) and recruitment of immune cells to the damaged muscle (Figure 14). DUX4-FL protein expression, apoptosis, and immune cell recruitment all peak at MD14, at which point there is also an increase in eMyHC positive, newly regenerated fibers as wells as fibrosis (Figures 16 and S8). Muscles at this stage produce ~60% of the force produced by ACTA1-MCM controls (Figure 5). Analysis of global differential gene expression supports the activation of apoptosis, the immune response, and the cell cycle, all three of which are much larger groupings than enriched in C2C12 cells expressing DUX4. This illustrates a key difference between performing studies *in vitro* using single cell types overexpressing DUX4 compared with studies of intact muscle expressing mosaic levels of DUX4 and containing all associated cell types (Figure 7).

A key component of the moderate model is that after MD14 both male and female mice recover on their own, regaining ~50% treadmill running by MD28, which coincides with decreases in apoptosis, eMyHC staining, and immune cell infiltration, although fibrosis remains. This is likely because DUX4-FL positive myofibers die and are being repaired and replaced using satellite cells that did not undergo transgene recombination, and thus have not activated DUX4 expression. Therefore, it is imperative to have the proper controls when using these models for preclinical testing of therapeutics. However, this also presents an opportunity whereby the mice can be re-injected with TMX at some point during treatment and progressive decline of the model can be assessed over several months. Preliminary experiments in our lab suggest that this is a viable possibility that needs further investigation and characterization. In addition, TMX can be adjusted to single higher-dose injections or multiple lower-dose injections to refine the model to meet investigational needs. The flexibility and tunability of this model is almost endless.

### Severe FSHD-like mouse model

While the moderate model can be assayed over the course of a month, the severe model is only useful for short-term analyses, as the DUX4-induced pathology is so severe that the mice require sacrifice no later than SD10. These mice do not show any signs of recovery. As in the moderate model, there are consistently different levels of transgene recombination and DUX4-fl expression among different anatomical muscles (Figure 1B). However, different muscles show the same patterns as in the moderate model (e.g., soleus is highest and quadriceps is lowest for both models), supporting the idea that accessibility to the TMX that is responsible for the variability. In addition, as seen for both the mild and moderate models, females and males of the severe model showed sex-specific differences, with females again being more severely affected. Upon induction of DUX4-fl expression, apoptosis (Figure 13), eMyHC expression (Figure 14), immune cell recruitment (Figure 15), and fibrosis (Figures 16 and S8) appear by SD6 and are all at the highest levels for any model at SD9. Conversely, *Mstn* levels are at their lowest for any model at SD9 (Figure S7). Treadmill stamina is significantly affected by SD6 and mice are immobile by SD9, with muscles producing ~25% of the force generated by ACTA1-MCM controls and ~50% of the force produced in the moderate model (Figures 4, 5, and S5). These markers for pathology are supported by global differential gene expression profiles that show greater enrichment for induced genes relating to apoptosis, immune response, and cell cycle, compared to the moderate model, while many muscle biology genes are significantly decreased (Figure 7).

## Conclusions

The goal in this study was to generate three levels of FSHD-like severity using our *FLExDUX4* mouse model and characterize the progression of pathology in ways useful to those performing preclinical testing of candidate DUX4-targeted therapeutics. We have provided an initial molecular, phenotypic, physiological, and histological characterization of three severity levels of inducible FSHD-like model mice. A key feature of these models is that the *ACTA1-MCM;FLExD* bi-transgenic mice express chronic low-level mosaic and skeletal muscle-specific DUX4-fl mRNA and protein. Thus, the moderate and severe models are not introducing DUX4 expression to a naive system, instead providing a situation similar to the bursts of DUX4 expression seen in FSHD myocytes [19]. It is well-documented that cells from asymptomatic FSHD subjects express DUX4, and cells from relatively healthy muscle biopsies from clinically affected FSHD patients express significant levels of DUX4 [9, 16, 17]. Thus, these models are recapitulating how we envision the DUX4 situation in FSHD, whereby genetically FSHD patients express a low level of DUX4 in their skeletal muscles and at some point, DUX4 levels significantly increase, along with muscle pathology. These bi-transgenic FSHD-like models allow investigators to recapitulate the chronic, low-levels of DUX4, followed by an investigator-controlled increase in DUX4 expression, and accompanying pathology, to whatever degree is desired.

Overall, these dose-dependent DUX4-fl FSHD-like phenotypic mouse models strongly support the DUX4 misexpression model for mediating FSHD pathogenesis [15, 18], and provide a useful and highly flexible tool for performing FSHD preclinical testing of therapeutic approaches targeting DUX4-fl mRNA and protein. Importantly for future analyses, we have shown sex-specific differences, anatomical muscle-specific differences, and model-specific differences that must be taken into account when using these FSHD-like mice. Within a single mouse, one can assess differentially affected muscles. Studying both sexes from a cross provides more fine-tuning of effects as well, with females being slightly but significantly more affected than the males. This provides even greater flexibility and utility for the model as a tool for studying FSHD and testing potential therapeutic approaches.

## Declarations

### Ethics approval

All animal procedures were approved by the local IACUC committee at the University of Nevada, Reno (#0701).

### Consent for publication

Not Applicable

### Availability of data and materials

Most data generated and analyzed during this study are included in the manuscript and supplemental data. Any data not included is available from the corresponding authors upon request. Raw RNA-seq data generated in this study has been deposited in the NCBI Gene Expression Omnibus (GEO) database under accession number (pending). Processed gene expression data has been deposited into GEO under accession number (pending).

### Authors’ contributions

TJ conceived of the study, performed experiments, analyzed data and wrote the manuscript. GW analyzed data and wrote the manuscript. PB performed experiments and analyzed data. SS performed experiments and analyzed data. MR performed experiments and analyzed data. RW performed experiments, analyzed data and wrote the manuscript. DB conceived of the study and wrote the manuscript. RB analyzed data and wrote the manuscript. PJ conceived of the study, analyzed data and wrote the manuscript. All authors read and approved the final manuscript.

## Acknowledgements

We thank Jennifer Burgess, Chris Carrino, Mick Hitchcock, PhD, Chris Hughes, and Daniel P. Perez for supporting our FSHD mouse work. We thank Dr. Charis Himeda for helpful discussions and editing the manuscript. This work was funded by grants from the Chris Carrino Foundation for FSHD, the FSH Society (FSHS-22012-01), the Muscular Dystrophy Association (MDA383364), and the National Institute of Arthritis and Musculoskeletal and Skin Diseases (R01 AR070432 and R21 AR070438) to PLJ, and from the National Institute of Neurological Disorders and Stroke P01 NS069539 to RKB. PLJ is supported by the Mick Hitchcock, PhD Endowed Chair of Medical Biochemistry at UNR Med. RKB is a Scholar of The Leukemia & Lymphoma Society.

## Abbreviations

DPI: days post-injection
EDL: extensor digitorum longus
FLExD: FLExDUX4
FSHD: facioscapulohumeral muscular dystrophy
GA: gastrocnemius
H&E: hematoxylin & eosin
IF: immunofluorescence
IP: intraperitoneal
MCM: MerCreMer
MD: moderate model day
Mstn: Myostatin
nLacZ: nuclear ß-galactosidase
PCR: polymerase chain reaction
qRT-PCR: quantitative reverse transcriptase PCR
RNA-seq: RNA sequencing
QUA: quadriceps
SD: severe model day
SOL: soleus
TA: tibialis anterior
TMX: tamoxifen
TUNEL: terminal deoxynucleotide transferase dUTP nick end labeling

## Supporting Information

**Figure S1: Mosaic tamoxifen dose-dependent recombination in gastrocnemius muscle of *ACTA1-MCM;R26^NZG^* bi-transgenic mice.** A) Cartoon depicting the cre-induced nuclear ß-galactosidase (nLacZ) expression from the *R26^NZG^* reporter mice. *ACTA1-MCM* mice express MerCreMer, a tamoxifen (TMX)-inducible Cre DNA recombinase under control of the skeletal muscle specific human skeletal actin gene (*ACTA1*) promoter. In *ACTA1-MCM; R26^NZG^* bi-transgenic mice, MerCreMer protein translocate to nucleus upon TMX binding and the resulting recombination causes the deletion of the loxP-flanked PGK-Neo cassette and expression of a nuclear localized ß-galactosidase protein, which is detected by X-Gal (blue) staining. B-E) X-Gal staining of cross sections of gastrocnemius muscles. B) The negative control *R26^NZG^/+* mice injected with tamoxifen. C) *ACTA1-MCM; R26^NZG^* mice have a low mosaic pattern of X-Gal signal in the absence of TMX induction indicating a very low percentage of cells have leaky MerCreMer protein translocation to the nucleus. *ACTA1-MCM; R26^NZG^* mice with one intraperitoneal (IP) injection of 5 mg/kg TMX (D) or two IP injections of 10 mg/kg TMX (E) show dose-dependent increases in recombined nuclei expressing nLacZ. All sections were stained with X-Gal for 50 minutes. Scale bar = 50 μm.

**Figure S2: Increased TMX dosage leads to increased mosaic recombination in skeletal muscle of *R26^NZG^, oACTA1-MCM* bi-transgenic mice.** Inverted images of X-Gal signal (white) overlaid with the DAPI signal (blue) were used to identify the nuclear recombination frequency in gastrocnemius muscle sections, described in Figure S1. A) The negative control ACTA1-MCM mice have no LacZ expression. B) Bi-transgenic ACTA1-MCM; R26^NZG^ mice in the absence of TMX show a low mosaic pattern of nuclear LacZ expression (231.5 X-Gal positive nuclei/mm^2^). C) Low level and D) high dose of TMX show 668.5 and 941.9 X-Gal positive nuclei/mm^2^, respectively. Scale bar = 100 μm

**Figure S3: The moderate and severe FSHD-like mouse models show significant weight loss.** Mice were weighed during time courses of TMX treatment prior to treadmill running (Figure 4). A) The moderately affected female mice (red line) were assayed prior to TMX injection (D0) and 3, 6, 10, 16, 23 and 29 days post-injection (DPI) and compared with age-matched female bi-transgenic mice (green line) and female *ACTA1-MCM* mice injected with TMX (blue line). B) The moderately affected male mice (red line) were assayed prior to TMX injection (D0) and 2, 6, 10, 14, 17, 21, 24 and 28 DPI and compared with age-matched male bi-transgenic mice (green line) and male *ACTA1-MCM* mice injected with TMX (blue line). C) Severely affected female mice (red line) were assayed prior to TMX injection (D0) and 2, 5, 7, and 8 DPI and compared with age-matched female bi-transgenic mice (green line) and *ACTA1-MCM* mice injected with TMX (blue line). D) Severely affected male mice (red line) were assayed prior to TMX injection (D0) and 2, 5, 7, and 8 DPI and compared with age-matched male bi-transgenic mice (green line) and *ACTA1-MCM* mice injected with TMX (blue line). The numbers of mice (n) for each experiment are same as in Figure 4. Significance = * p<0.05, ** p<0.01, ***p<.001, calculated between bi-transgenic + TMX and bi-transgenic no TMX (green) or *ACTA1-MCM* + TMX (blue).

**Figure S4: *Ex vivo* analysis of muscle function shows significant muscle weakness in the female mild and <u>moderately</u> affected FSHD-like mouse models.** EDL muscles from female *ACTA1-MCM* +TMX (*MCM* +TMX, control, blue, n=9) mice, bi-transgenic mild mice without TMX (*MCM;FLExD* mild, green, n=9), bi-transgenic +TMX moderate mice (*MCM;FLExD* Mod, yellow, n=6), and bi-transgenic +TMX severe mice (*MCM;FLExD* Severe, red, n=3) were isolated at MD14 or SD10 and assayed for A) maximum twitch, B) maximum tetanus, C) force frequency, and then normalized to muscle cross sectional area to provide specific force measurements (Figure 5A-C). Significance = * p<0.05, ** p<0.01, ***p<.001; blue asterisks indicate significance compared with *MCM* +TMX control mice, green asterisks indicate significance when compared with the bi-transgenic *MCM;FLExD* mild model mice, and yellow asterisks indicate significance when compared with the bi-transgenic *MCM;FLExD* moderate mice.

**Figure S5: *Ex vivo* analysis of muscle function shows significant muscle weakness in the male mild and severely affected FSHD-like mouse models.** EDL muscles from male *ACTA1-MCM* +TMX (MCM +TMX, control, blue, n=5) mice, bi-transgenic mice without TMX (*MCM;FLExD*, mild model, green, n=9), and bi-transgenic mice +TMX (*MCM;FLExD* +TMX, severe model, red, n=11) were assayed for A) maximum twitch force, B) maximum tetanus force, C) force frequency, and then normalized for muscle cross sectional area to provide D) specific maximum twitch force, E) specific maximum tetanus force, and F) specific force frequency. Significance = * p<0.05, ** p<0.01, ***p<.001; blue asterisks indicate significance compared with the *ACTA1-MCM* control mice and green asterisks indicate significance compared with bi-transgenic *MCM;FLExD* mild model mice.

**Figure S6: The heart is not affected by TMX treatment in control or bi-transgenic animals.** H&E staining of cryosections of heart isolated from A) *ACTA1-MCM* mice, and bi-transgenic *ACTA1-MCM;FLExD* mice B) without TMX and C) with the high-dose (2X 10 mg/kg) of TMX. Scale bar = 100 μm

**Figure S7: *Mstn* gene expression decreases with increased *DUX4-fl* expression in the FSHD-like mouse models.** *Mstn* mRNA levels from 3-month old mice were assayed by qRT-PCR, for the models as indicated. The moderate model was assayed at MD9 and the severe model at SD9. Levels were normalized to *Rpl37* mRNA levels. Significance was calculated using one-way Anova with uncorrected Fisher’s LSD. * p<0.05, ** p<0.01, *** p<0.001, ns= not significant

**Figure S8: Significant fibrosis in late stages of the moderate and severe FSHD-like mouse models.** Summary of Sirius red staining for tibialis anterior muscle sections for mild (green), moderate (yellow), and severe (red) FSHD-like mouse models compared with *ACTA1-MCM* control (blue). Data is plotted as percent fibrotic area. Significance was calculated using the least significant difference test, with *= p<.05, **= p<.01, and ***= p<.001.

**Figure S9: Cardiac muscle from FSHD-like mouse models shows no signs of increased fibrosis.** Hearts were isolated from the same mice assayed in Figure 14 for skeletal muscle fibrosis. Control *ACTA1-MCM* mice (blue), bi-transgenic mild model mice (green) and the severe model mice at SD9 (red) show no significant differences in heart fibrosis.

**Table S1: Significantly altered gene expression in FSHD-like models**

**Table S2: Differentially expressed genes**

**Table S3: Intersection of misregulated genes in models with FSHD patients**

**Table S4: GO term superterms**

**Table S5: GO superterm genes**

**Table S6: MISO alternative splicing analysis for SE and RI**

